# Mechanistic basis for protein conjugation in a diverged bacterial ubiquitination pathway

**DOI:** 10.1101/2024.11.21.623953

**Authors:** Qiaozhen Ye, Minheng Gong, Lydia R. Chambers, Kevin D. Corbett

## Abstract

Ubiquitination is a fundamental and highly conserved protein post-translational modification pathway, in which ubiquitin or a ubiquitin-like protein (Ubl) is typically conjugated to a lysine side chain of a target protein. Ubiquitination is a multistep process initiated by adenylation of the Ubl C-terminus, followed by sequential formation of 2-3 Ubl∼cysteine thioester intermediates with E1, E2, and E3 proteins before formation of the final Ubl-lysine isopeptide bond^1^. Ubiquitination is conserved across eukaryotes, and recent work has also revealed at least two related bacterial pathways that perform protein conjugation in the context of antiphage immunity^2–5^. Bioinformatics analysis has hinted at the existence of additional, as-yet uncharacterized, pathways in bacteria that could perform protein conjugation using ubiquitination-like machinery^6–8^. Here we describe the architecture and biochemical mechanisms of Bub (<u>b</u>acterial <u>ub</u>iquitination-like) pathways, revealing strong structural parallels along with striking mechanistic differences when compared to eukaryotic ubiquitination pathways. We show that Bub operons encode functional E1, E2, and Ubl proteins that are related to their eukaryotic counterparts but function entirely through oxyester, rather than thioester, intermediates. We also identify a novel family of serine proteases in Bub operons with a conserved serine-histidine catalytic dyad. The genomic context of Bub operons suggests that, like other bacterial ubiquitination-related pathways, they also function in antiphage immunity. Overall, our results reveal a new family of bacterial ubiquitination-related pathways with unprecedented biochemical mechanisms in both protein conjugation and deconjugation.

## Introduction

Ubiquitination and related pathways typically involve three proteins termed E1, E2, and E3 (ref. ^1^). E1 proteins initiate the reaction by adenylating the C-terminus of a Ubl, then the E1 catalytic cysteine attacks this intermediate to generate a Ubl∼E1 thioester (‘∼’ denotes a reactive covalent intermediate). The Ubl is next transferred to the E2 catalytic cysteine to form a second thioester intermediate. Finally, the Ubl becomes covalently attached to a target through the action of an E3 protein, typically forming an isopeptide linkage to a lysine side chain’s primary amine group. In eukaryotes, ubiquitination usually marks proteins for proteasome-mediated degradation, while other Ubl pathways regulate protein stability, conformation, and protein-protein interactions^9,10^. At least one Ubl pathway, the ISG15 pathway, functions in innate immunity by targeting viral proteins for modification by ISG15 (refs. ^11–13^).

Ubiquitination pathways are found throughout eukaryotes, but bacteria have historically not been considered to possess ubiquitination-like pathways that perform protein conjugation. Recently, comparative genomics of thousands of bacterial genomes identified several sparsely-distributed families of bacterial operons that encode different combinations of predicted Ubl, E1, E2, E3, and peptidase proteins (termed deubiquitinases or DUBs)^6–8^. Structural, biochemical, and functional analysis of two such operon families - termed Type II CBASS (<u>c</u>yclic-oligonucleotide-<u>b</u>ased <u>a</u>ntiphage <u>s</u>ignaling <u>s</u>ystem) and Type I/II Bil (<u>B</u>acterial ISG15-like protein) pathways -has demonstrated that they are evolutionarily related to eukaryotic ubiquitination pathways and mediate protein conjugation using a parallel biochemical mechanism^2–5^. Both Type II CBASS and Bil are immune pathways that protect their bacterial hosts against bacteriophage (phage) infection. In the case of Bil operons, whose Ubls show structural similarity to the eukaryotic Ubl ISG15, antiphage immunity arises from specific conjugation of its Ubl to a phage tail protein^5^.

Alongside Type II CBASS and Bil operons, other families of bacterial operons encoding putative ubiquitination-like pathways have been identified bioinformatically but not yet functionally characterized. Here we focus on an uncharacterized operon family previously termed “6E” or “DUF6527”, which encodes combinations of predicted E1, E2, peptidase, and Ubl proteins plus an unknown protein with a DUF6527/Pfam20137 domain^2,6,7^. 6E/DUF6527 operons’ E2-like proteins were previously noted to lack a conserved catalytic cysteine and have therefore been predicted to be inactive for Ubl transfer^6^. 6E/DUF6527 operons’ E1-like proteins also lack a conserved catalytic cysteine (see **Results**), raising further doubt as to whether these operons encode functional ubiquitination-like pathways.

Here, we show that 6E/DUF6527 operons, which we term “Bub” for <u>b</u>acterial <u>ub</u>iquitination-like operons, are *bona fide* protein conjugation pathways whose E1, E2, and Ubl proteins show structural and functional similarity to canonical eukaryotic ubiquitination machinery. Combining sequence analysis, high-resolution structures, and mechanistic biochemistry, we find that Bub operons function through oxyester, rather than thioester, intermediates. The Bub E1 adenylates its cognate Ubl and forms a Ubl∼tyrosine oxyester, and the Bub E2 forms a Ubl∼serine oxyester. Finally, we find that DUF6527/Pfam20137 proteins are a novel family of serine proteases, with a conserved serine-histidine catalytic dyad that can cleave both peptide and isopeptide bonds. Together, our data reveal the architecture and function of a highly diverged family of bacterial ubiquitination-like pathways, and highlight an unprecedented biochemical mechanism for Ubl conjugation.

## Results

### Bacterial Bub operons encode diverged ubiquitination-related pathways

Prior bioinformatics studies have defined several families of bacterial operons that encode proteins related to E1, E2, E3, and deubiquitinase/peptidase proteins^6–8,14^. A subset of these operon families also encodes a predicted ubiquitin-like (Ubl) protein, the first of which is termed Bil (<u>B</u>acterial ISG15-like). Bil operons have been shown to protect their host from phage infection by conjugating a Ubl to a phage structural protein to inhibit phage assembly and infectivity^5,15^. The two families of Bil operons, Type I and Type II, both encode a Ubl (BilA), an E2 (BilB), a JAB/JAMM-family peptidase (BilC), and an E1 protein encoding tandem inactive and active adenylation domains plus a mobile catalytic cysteine-containing domain (IAD-AAD-CYS; BilD)^15^. Bil operons have been shown to mediate Ubl-target conjugation, and to generate *bona fide* Ubl-target lysine isopeptide linkages^4,5^. Bil operons do not encode predicted E3 proteins; rather, their E2 proteins are proposed to directly recognize ubiquitination targets^4,5^.

Another major group of Ubl-containing operons has previously been termed “6E” or “DUF6527”, and here is termed Bub (<u>B</u>acterial <u>ub</u>iquitination-like)^2,6,7^. Bub operons can be divided into two families based on their gene complement: Type I Bub (previously 6E.1) operons encode a predicted Ubl-E2 fusion (BubAB), a JAB/JAMM family peptidase-E1 fusion (BubCD), and a DUF6527 protein (BubE; **Figure 1a, Table S1**). Type II Bub (previously 6E.2) operons encode a Ubl (BubA), an E2-E1 fusion (BubBD), and a DUF6527 protein (BubE), and typically do not encode a JAB/JAMM family peptidase. The Ubl proteins in both Type I and Type II Bub operons show high structural diversity, encoding 1-3 ubiquitin-like β-grasp domains and a variety of N-terminal domains^4,16^.

**Figure 1.**
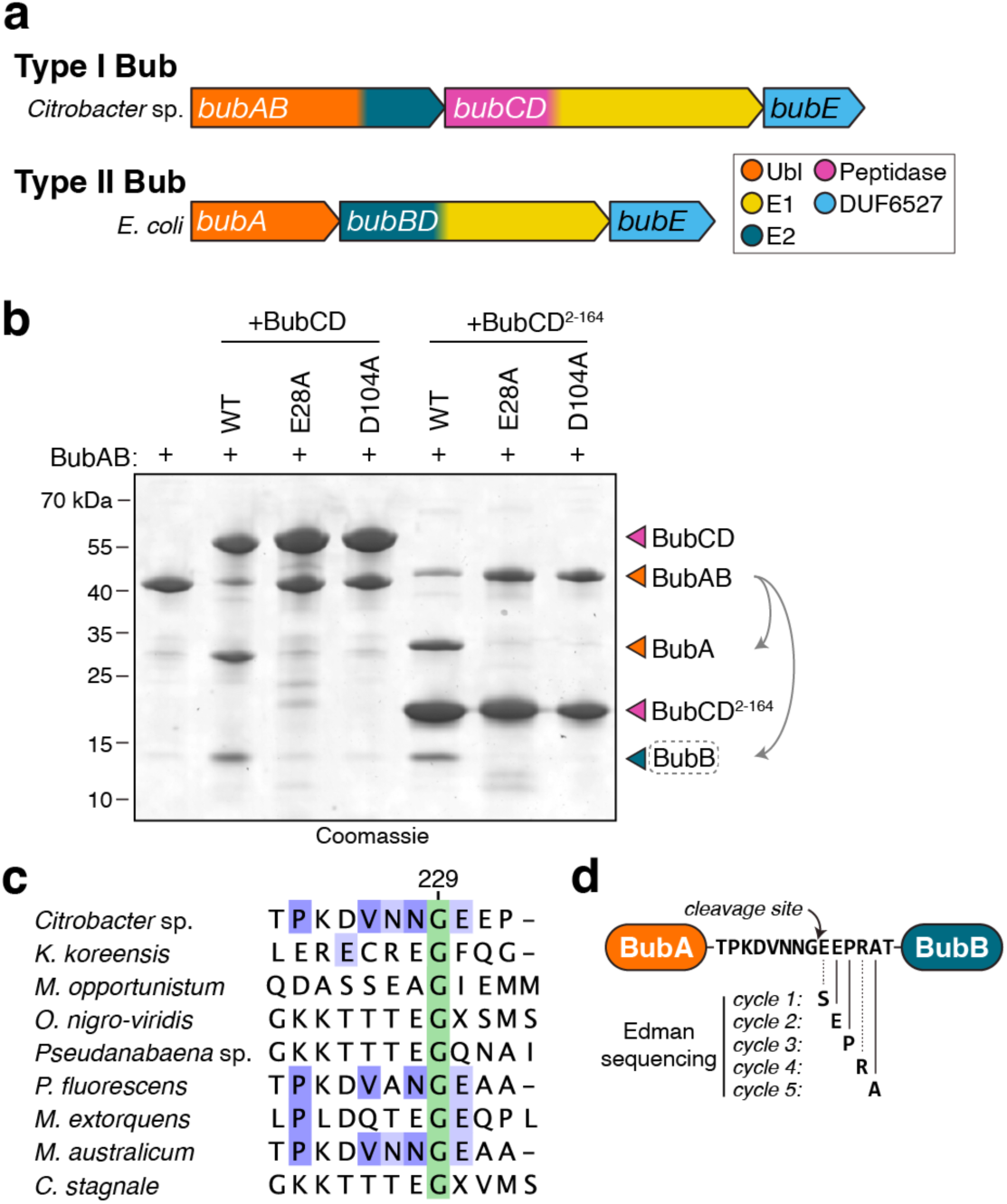
Identification of Bub operons. **(a)** Operon architecture of Type I and Type II Bub (<u>b</u>acterial <u>ub</u>iquitination-like) operons. See **Table S1** for a list of identified operons and accession numbers. **(b)** Purified *Citrobacter* BubCD or the isolated peptidase domain (residues 2-164) was incubated with BubAB and the result analyzed by SDS-PAGE and Coomassie blue staining. BubCD/BubCD^2–164^ constructs used were wild-type (WT), E28A, or D104A. The C-terminal cleavage product of BubAB cleavage (BubB; dotted box) was isolated and subjected to N-terminal sequencing by Edman degradation (panel d). **(c)** Sequence alignment of BubAB proteins (from an alignment of 107 BubAB proteins from Type I Bub operons) showing conservation of the C-terminal glycine residue of the BubA region (green; G229 in *Citrobacter* BubAB). Proteins shown are from *Citrobacter* sp. RHBSTW-00271 (IMG 2938140956), *Kangiella koreensis* SW-125 (IMG 644987647), *Mesorhizobium opportunistum* WSM2075 (IMG 2503204321), *Oscillatoria nigro-viridis* PCC 7112 (IMG 2504086278), *Pseudanabaena* sp. PCC 7367 (IMG 2504678157), *Pseudomonas fluorescens* NZ011 (IMG 2506358179), *Methylorubrum extorquens* DSM 13060 (IMG 2507323553), *Mesorhizobium australicum* WSM2073 (IMG 2509393842), and *Cylindrospermum stagnale* PCC 7417 (IMG 2509772370) (see **Table S1**). **(d)** N-terminal sequencing of the C-terminal cleavage product of BubAB (dotted box in panel B) by Edman degradation. Dotted lines (cycles 1 and 4) indicate low-confidence amino acid calls. See **Figure S1** for detailed data.

During manual inspection of identified Bub operons, we noticed that a large fraction are associated with transcriptional repressors previously shown to be associated with diverse antiphage immune pathways^17,18^. Out of 544 total Bub operons, 43 (8%) are associated with an HTH-WYL transcription factor related to CapW/BrxR, and 168 (31%) are associated with a CapH+CapP gene pair (**Table S1**). Both CapW/BrxR and CapH+CapP have been shown to drive expression of their associated immune pathway in response to phage infection and/or DNA damage^17,18^, and the association of these regulators with Bub operons suggests that they similarly function in antiphage immunity or stress response.

### BubCD cleaves BubAB to generate a functional ubiquitin-like protein

To determine the mechanistic basis for Bub operon function, we expressed and purified BubAB, BubCD, and BubE from a Type I Bub operon from *Citrobacter* sp. RHBSTW-00271 (**Figure 1a, Table S2**). As in all Type I Bub operons, this operon’s Ubl protein BubA is encoded as a fusion to the predicted E2 BubB. To determine whether BubAB is proteolytically processed to generate a standalone BubA capable of conjugation to target proteins, we incubated BubCD or its N-terminal JAB/JAMM-family peptidase domain (residues 2-164) with purified BubAB. In the presence of wild-type BubCD, but not mutants in which the predicted active site residues E28 or D104 were mutated to alanine, we observed cleavage of BubAB into two fragments (**Figure 1b, Table S2**). We subjected the cleaved product to N-terminal sequencing by Edman degradation, and found that BubCD cleaves BubAB immediately C-terminal to the conserved glycine G229 (**Figure S1**, **Figure 1c-d**). Thus, the N-terminal peptidase domain of BubCD cleaves the BubAB pre-protein to generate a Ubl with an exposed C-terminal glycine. For the remainder of our experiments, we separately expressed BubA (BubAB residues 1-229) and BubB (BubAB residues 230-362).

### BubCD is a homodimeric peptidase-E1 fusion lacking a catalytic cysteine

We next determined a 1.8 Å resolution structure of wild-type *Citrobacter* BubCD and a 2.0 Å resolution structure of a D104A peptidase domain mutant (BubCD^D104A^; **Table S3**). BubCD adopts a two-domain architecture with an N-terminal JAB/JAMM family peptidase domain rigidly docked against a C-terminal E1-like adenylation domain (**Figure 2a-b**). In both structures, BubCD forms a homodimer via its E1 domains (**Figure 2b**). Structurally, the BubCD E1 domain most closely resembles other homodimeric E1/E1-like proteins, including human UBA5, a minimal homodimeric E1 (4.9 Å r.m.s.d. over 168 Cα atoms), and the bacterial E1-like protein ThiF (4.4 Å r.m.s.d. over 224 Cα atoms; **Figure S2a-c**). BubCD possesses all necessary residues for the reaction of ATP with a Ubl C-terminus to generate an adenylated intermediate (**Figure 2c**).

**Figure 2.**
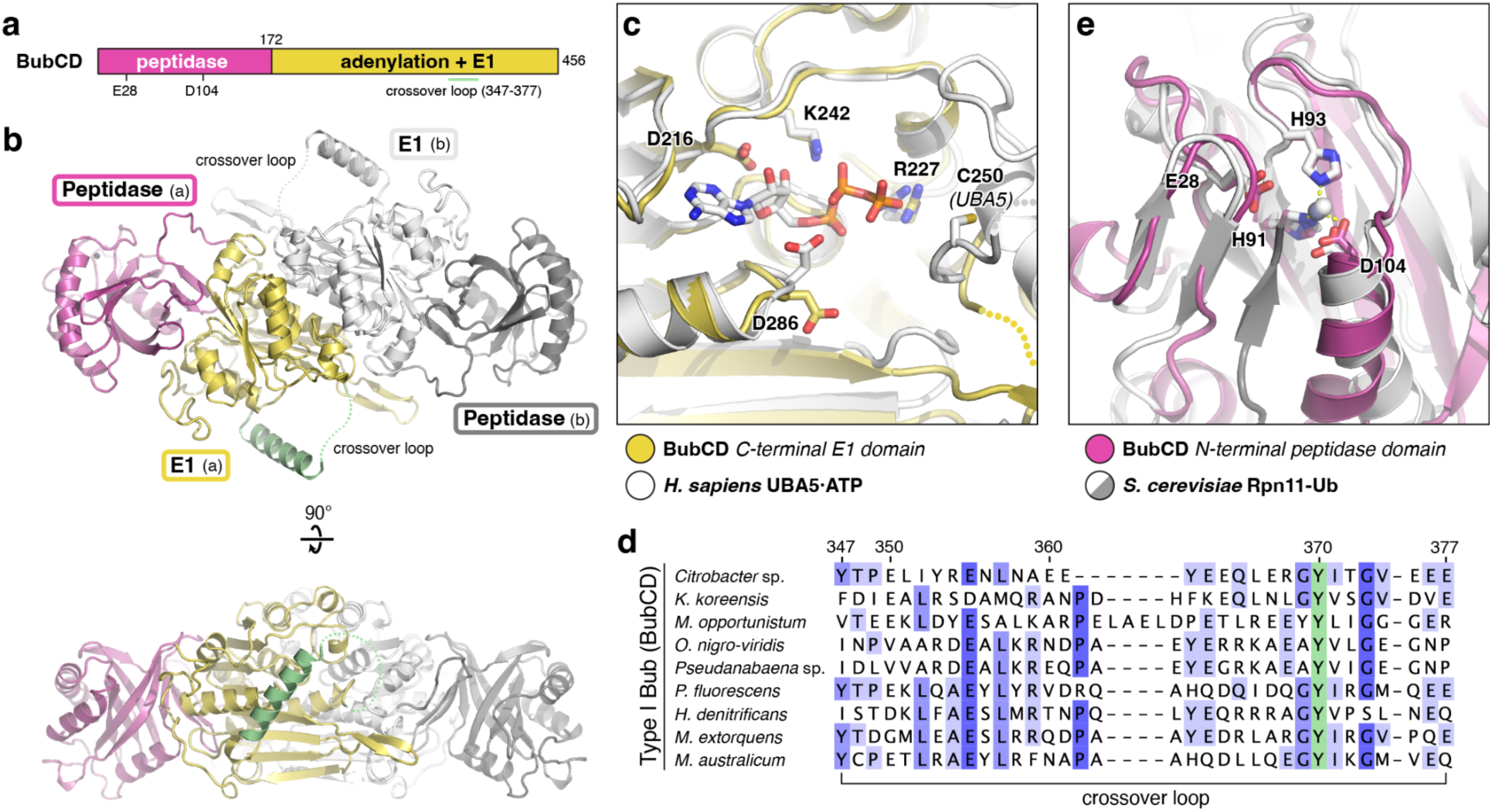
Structure of BubCD. **(a)** Schematic of *Citrobacter* BubCD, showing the N-terminal BubC peptidase domain (magenta; putative active site residues indicated) and the C-terminal BubD adenylation + E1 domain (yellow; crossover loop marked with green line). **(b)** Two views of the BubCD dimer, with one protomer colored as in panel (a) and the other colored gray (BubC peptidase) and white (BubD adenylation + E1). The crossover loop of one protomer is colored light green, and the disordered portion of this loop (residues 367-376) is indicated by a dotted line. **(c)** Closeup of the adenylation active site of BubCD (yellow) overlaid with that of *H. sapiens* UBA5 bound to ATP (white; PDB ID 6H78)^19^. Conserved ATP-binding active site residues are shown as sticks and labeled (BubCD numbering). The catalytic cysteine of UBA5 (C250) is labeled. **(d)** Sequence alignment of the crossover loop of BubCD proteins in Type I Bub operons (from an alignment of 137 unique BubCD sequences; see **Table S1**). The conserved tyrosine Y370 is highlighted in green; blue highlights show the level of conservation in the full alignment. Proteins shown are from *Citrobacter* sp. RHBSTW-00271 (IMG 2938140955), *Kangiella koreensis* SW-125 (IMG 644987648), *Mesorhizobium opportunistum* WSM2075 (IMG 2503204320), *Oscillatoria nigro-viridis* PCC 7112 (IMG 2504086279), *Pseudanabaena* sp. PCC 7367 (IMG 2504678156), *Pseudomonas fluorescens* NZ011 (IMG 2506358178), *Hyphomicrobium denitrificans* 1NES1 (IMG 2507044886), *Methylorubrum extorquens* DSM 13060 (IMG 2507323552), and *Mesorhizobium australicum* WSM2073 (IMG 2509393843). **(e)** Closeup of the peptidase active site of BubCD (magenta) overlaid with that of *S. cerevisiae* Rpn11 (white) bound to ubiquitin (gray; PDB ID 5U4P)^20^; bound Zn^2+^ ions are shown as spheres. Conserved active site residues are shown as sticks and labeled (BubCD numbering).

All known E1 proteins possess a conserved cysteine residue either in the adenylation domain’s “crossover loop” or in an α-helical domain inserted into the crossover loop, which attacks the adenylated Ubl to form a Ubl∼cysteine thioester intermediate^1^. For example, human UBA5 contains a catalytic cysteine, C250, near the C-terminus of its crossover loop (**Figure 2c**)^19^. Curiously, BubCD’s crossover loop (residues 347-377) does not contain a cysteine, and sequence alignments show that BubCD proteins generally lack cysteine residues in their crossover loops. Indeed, the only invariant residue in the crossover loop of BubCD is a tyrosine, Y370 (**Figure 2d**). While a tyrosine hydroxyl group could theoretically form an oxyester linkage with a Ubl C-terminus, to date Ubl∼tyrosine oxyester linkages have not been experimentally demonstrated, either as a reaction intermediate or as a final product of Ubl conjugation.

The N-terminal domain of BubCD resembles other JAB/JAMM family peptidase proteins, including the *S. cerevisiae* proteasome subunit Rpn11 (5.4 Å r.m.s.d. over 144 Cα atoms)^20^ and an archaeal peptidase, *Pyrococcus furiosus* JAMM1 (3.8 Å r.m.s.d. over 128 Cα atoms)^21^. In the structure of wild-type BubCD, a Zn^2+^ ion in the active site is coordinated by conserved active site residues H91, H93, and D104, with the catalytic glutamate residue E28 nearby (**Figure 2e**). In the structure of the BubCD^D104A^ mutant, we find that Zn^2+^ binding is lost and the active site is remodeled, with residues 93-108 adopting a different structure than in wild-type BubCD. This rearrangement destroys the substrate-binding pocket (**Figure S2d**), rendering this domain unable to bind BubA.

### Structures of BubCD separately bound to BubA and BubB

Next, we separately reconstituted complexes of BubCD^D104A^ with BubA and BubB. Because *Citrobacter* BubA possesses three β-grasp domains and forms helical filaments via domains 1 and 2 (ref. ^16^), we used a truncated BubA construct encoding only β-grasp domain 3 (residues 155-229). We determined a 2.3 Å resolution structure of BubCD^D104A^ bound to BubA^155–229^, which showed BubA binding to the BubCD adenylation domain in a manner consistent with other E1/E1-like proteins bound to their respective Ubls, including *H. sapiens* UBA5-UFM1 and *E. coli* ThiF-ThiS (**Figure 3a-c, Figure S3a-c**). In BubCD, an extended β-hairpin (residues 312-328) extends outward from the adenylation domain to buttress the bound BubA (**Figure 3c**). The BubA C-terminal glycine 229 is positioned for adenylation, and the BubCD crossover loop is ordered, forming two short α-helices followed by an extended random-coil segment. In this conformation, BubCD Y370 is pointed away from the enzyme’s adenylation active site (**Figure 3c**).

**Figure 3.**
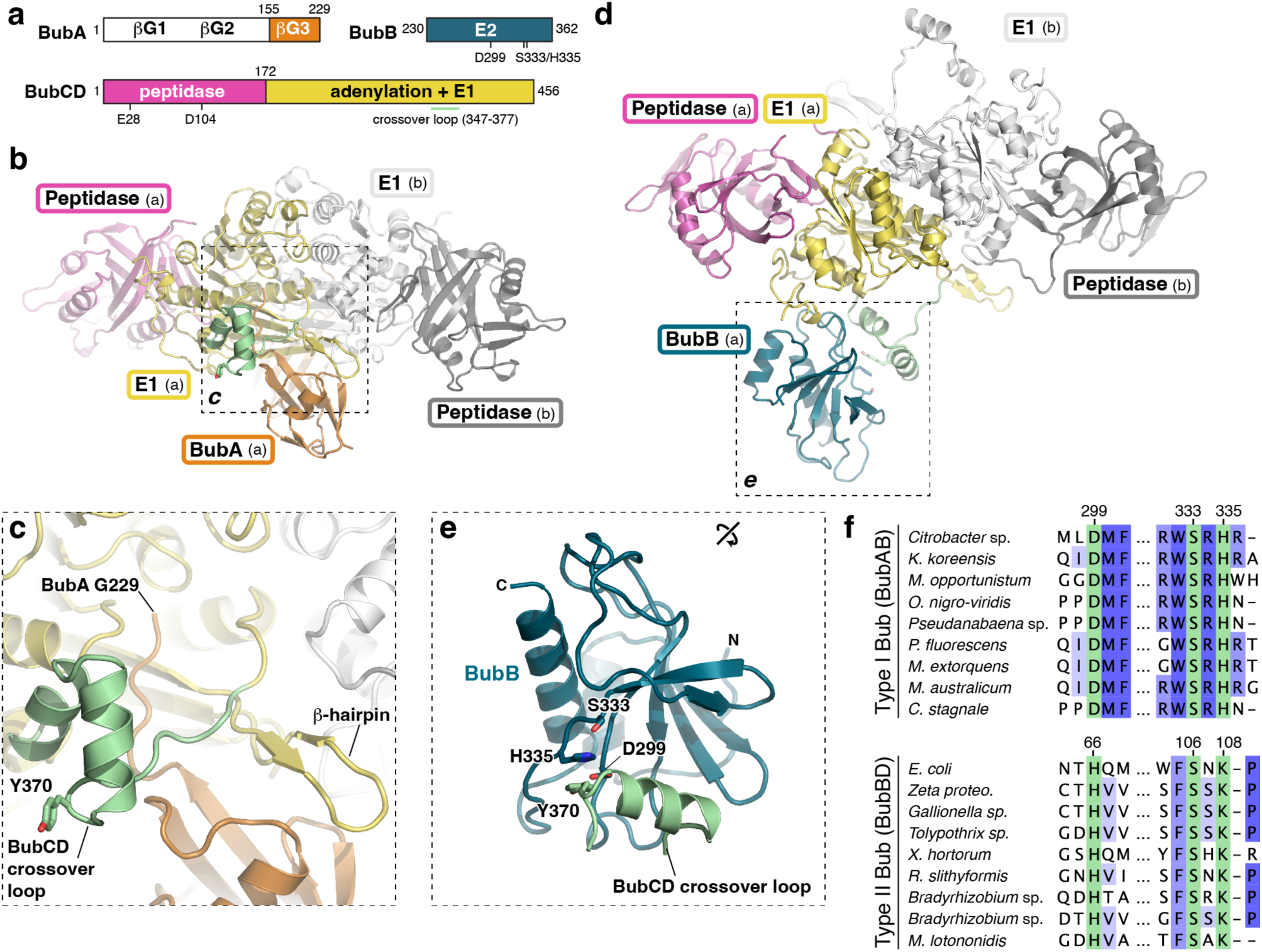
Structures of BubCD bound to BubA and BubB. **(a)** *Top:* Schematic of *Citrobacter* BubA and BubB (after processing by BubCD), with the three β-grasp domains of BubA marked βG1-βG3, and the putative active site residues S333 and H335 in the BubB E2 domain marked. *Bottom:* Schematic of *Citrobacter* BubCD. **(b)** Structure of the *Citrobacter* BubCD-BubA (155-229) complex, with one BubCD protomer colored as in panel (a) and the opposite protomer colored white/gray. **(c)** Close-up of the adenylation active site of BubCD, showing the C-terminus of BubA (G229) in the BubCD active site, and the BubCD crossover loop (light green) with conserved tyrosine Y370 shown as sticks. **(d)** Structure of the *Citrobacter* BubCD-BubB complex, with one BubCD protomer colored as in panel (a) and the opposite protomer colored white/gray. **(e)** Closeup of BubB (dark blue) interacting with the BubCD crossover loop (light green), showing the close proximity of putative BubB active site residues S333 and H335 to the conserved BubCD tyrosine Y370. **(f)** Sequence alignment of the putative active site residues of BubB in Type I Bub (*top*, from an alignment of 107 unique BubAB proteins, see **Table S1**) and Type II Bub (*bottom*, from an alignment of 400 unique BubBD proteins, see **Table S1**). Conserved putative active site residues are highlighted in green; blue highlights show the level of conservation in the full alignment. BubAB proteins shown are from *Citrobacter* sp. RHBSTW-00271 (IMG 2938140956; numbering shown), *Kangiella koreensis* SW-125 (IMG 644987647), *Mesorhizobium opportunistum* WSM2075 (IMG 2503204321), *Oscillatoria nigro-viridis* PCC 7112 (IMG 2504086278), *Pseudanabaena* sp. PCC 7367 (IMG 2504678157), *Pseudomonas fluorescens* NZ011 (IMG 2506358179), *Methylorubrum extorquens* DSM 13060 (IMG 2507323553), *Mesorhizobium australicum* WSM2073 (IMG 2509393842), and *Cylindrospermum stagnale* PCC 7417 (IMG 2509772370). BubBD proteins shown are from *E. coli* ZDHYS365 (NCBI WP_063617865.1; numbering shown), Zeta proteobacterium SCGC AB-604-B04 (IMG 2264879632), *Gallionella* sp. SCGC AAA018-N21 (IMG 2264885673), *Tolypothrix* sp. PCC 7601 (IMG 2501546803), *Xanthomonas hortorum pv. gardneri* 7 (IMG 2501866950), *Runella slithyformis* LSU4, DSM 19594 (IMG 2505789128), *Bradyrhizobium* sp. WSM1417 (IMG 2507506257), *Bradyrhizobium* sp. WSM471 (IMG 2508545363), and *Microvirga lotononidis* WSM3557 (IMG 2509076069).

Next, we determined a 2.5 Å resolution structure of BubCD^D104A^ bound to BubB (**Table S3**). In the structure, only one protomer of the BubCD dimer is bound to BubB, creating a complex with 2:1 stoichiometry (**Figure 3d**). BubB adopts a diverged UBC fold that most closely resembles the E2 domain of the bacterial E2-E1 protein Cap2 (4.3 Å r.m.s.d. over 120 Cα atoms) and *H. sapiens* UBE2D2 (3.8 Å r.m.s.d. over 104 Cα atoms; **Figure S3d-f**). Like the Cap2 E2 domain and other non-canonical E2s, BubB lacks a characteristic pair of α-helices at the C-terminus of the UBC fold (**Figure S3d-f**). Like BubCD, BubB also apparently lacks a catalytic cysteine. Instead, BubB proteins possess an invariant serine (S333 in *Citrobacter* BubB) in a structurally-equivalent position to other E2s’ catalytic cysteines (**Figure 3e, Figure S3d-f**). In Type I Bub operons with BubAB fusion proteins (including *Citrobacter* Bub), the invariant serine is accompanied by a nearby invariant histidine (H335 in *Citrobacter* BubB) and a nearby invariant aspartate (D299 in *Citrobacter* BubB; **Figure 3e-f**). In Type II Bub operons with BubBD fusion proteins, the invariant serine is accompanied by invariant lysine and histidine residues (**Figure 3f**). These accompanying residues (histidine+aspartate in Type I Bub, lysine+histidine in Type II Bub) may alter the chemical environment of the BubB active site to increase the reactivity of the nearby serine, making it more amenable to forming a Ubl∼serine oxyester intermediate and then conjugating the Ubl to a target. In our structure of *Citrobacter* BubCD^D104A^ bound to BubB, BubCD Y370 is positioned close to the invariant serine-histidine residue pair in BubB (**Figure 3d-e**), potentially mimicking the conformation of a Ubl handoff from BubCD Y370 to BubB S333.

### Type I Bub operons function through oxyester intermediates

To visualize a ternary complex of BubCD, BubA, and BubB, we separately purified *Citrobacter* BubCD (D104A/Y370F double mutant), BubA^155–229^, and BubB, crystallized the complex in the presence of Mg^2+^ and ATP, and determined a 1.93 Å resolution structure (**Table S3**). In the resulting structure, the asymmetric unit contains a dimer of BubCD, two protomers of BubB, and one protomer of BubA^155–229^ (**Figure 4a**). Each protomer of BubCD is bound to BubB in a manner equivalent to our structure of BubCD^D104A^-BubB, with the BubCD crossover loop ordered and the conserved Y370 (mutated to phenylalanine) juxtaposed to BubB residues D299, S333, and H335 (**Figure 4b-c**). Within the crystal, one BubB protomer clashes with its symmetry mate through crystal packing; we therefore refined this protomer at 50% occupancy (see **Materials and Methods**). Within the BubCD dimer, we observed that one BubCD protomer is bound to BubA^155–229^ equivalently to our structure of the BubCD^D104A^-BubA^155–229^ complex and does not contain a nucleotide in the BubCD adenylation active site (**Figure 4b**). The second BubCD protomer, meanwhile, is not bound to BubA^155–229^ but contains a well-ordered AMP molecule in its adenylation active site (**Figure 4c**). Overall, this structure confirms that BubCD can interact with both BubA and BubB simultaneously, and that BubCD can bind a nucleotide in its adenylation active site.

**Figure 4.**
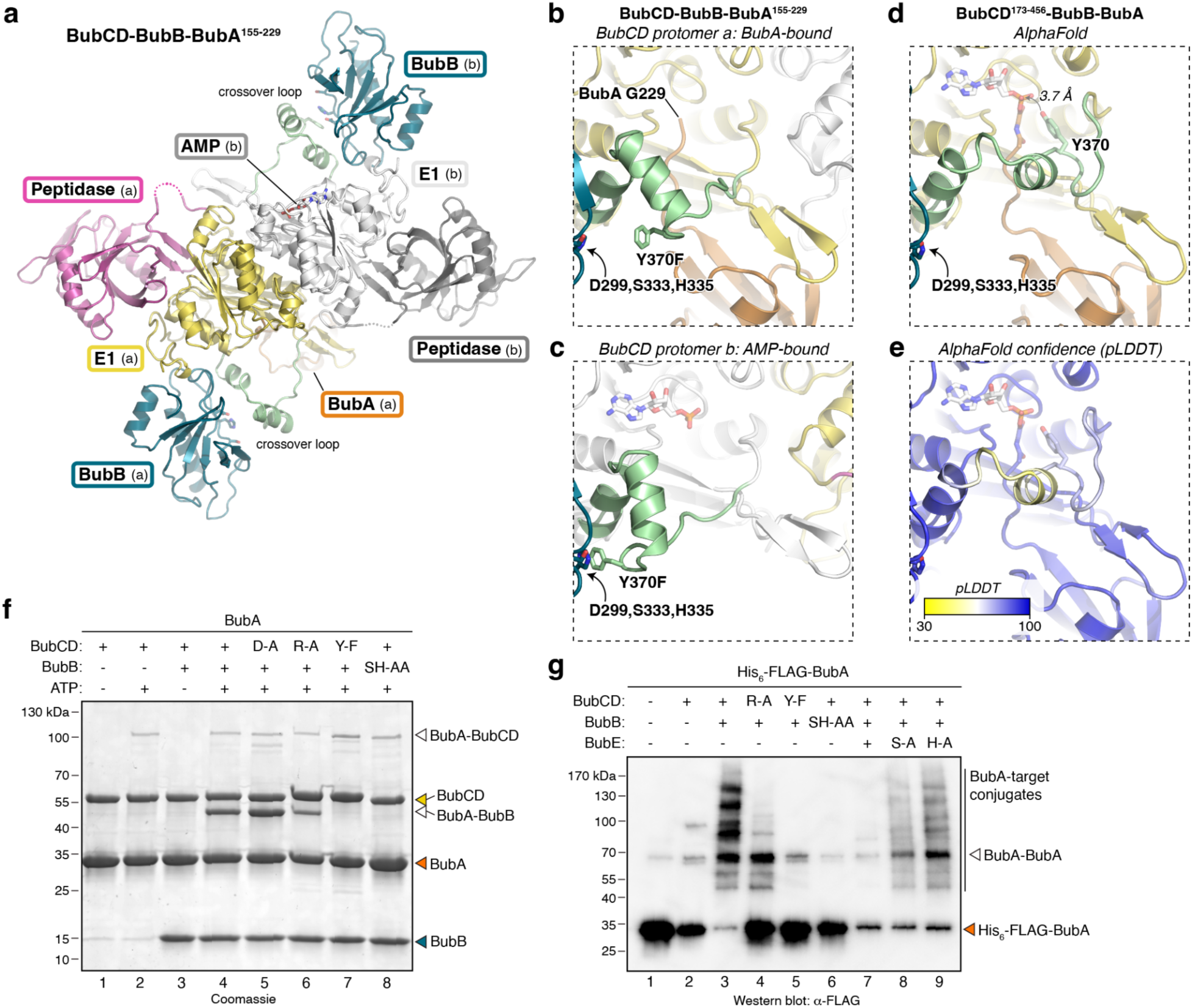
Structure and activity of *Citrobacter* BubCD-BubB-BubA. **(a)** Structure of BubCD (D104A/Y370F double mutant) bound to BubB and BubA^155–229^. The complex is asymmetric, with BubCD protomer (a) bound to BubB and BubA^155–229^, and BubCD protomer (b) bound to BubB and AMP (shown as sticks). **(b-c)** Closeup view of the BubCD adenylation active site in protomer (a) (panel b) and protomer (b) (panel c). BubCD Y370 (mutated to phenylalanine) and BubB S333 and H335 are shown as sticks and labeled. **(d)** View equivalent to panels (b) and (c), showing an AlphaFold 2 model of BubCD (residues 173-456) bound to BubB and BubA (see **Figure S4a-b**) with AMP manually modeled based on the AMP-bound BubCD structure. This model shows BubCD Y370 3.7 Å away from the AMP α-phosphate, primed for attack and formation of an oxyester intermediate. **(e)** View equivalent to panel (d) showing AlphaFold 2 confidence (pLDDT), colored from yellow (pLDDT values 30 or lower; low confidence) to blue (pLDDT value 100; high confidence). The pLDDT value for BubCD Y370 is 74.88. **(f)** SDS-PAGE analysis of *in vitro* reactions with *Citrobacter* BubA, BubB, and BubCD. BubCD D-A: D104A peptidase domain mutant; R-A: R227A adenylation domain mutant; Y-F: Y370F crossover loop mutant. BubB SH-AA: S333A/H335A double mutant. **(g)** Anti-FLAG western blot from co-expression and denatured Ni^2+^-affinity purification of His_6_-FLAG-tagged *Citrobacter* BubA with BubCD, BubB, and BubE (wild-type and indicated mutants). BubCD R-A: R227A adenylation domain mutant; Y-F: Y370F crossover loop mutant. BubB SH-AA: S333A/H335A double mutant. BubE S-A: S92A; H-A: H102A.

Our structural data and sequence alignments suggest that BubCD Y370 could play a catalytic role, but our structures do not reveal how this residue could react with a putative adenylated BubA intermediate. We performed a series of structure predictions using AlphaFold^22,23^ with different combinations of BubCD, BubA, and BubB to explore potential conformations of BubCD that were not captured in our crystal structures. When we used AlphaFold 2 to predict the structure of *Citrobacter* BubCD E1 domain (residues 173-456) bound to BubA and BubB, the resulting prediction showed BubA and BubB bound to BubCD exactly as in our crystal structures (**Figure S4a-b**). The only major difference between the predicted structure and our crystal structures was the conformation of the BubCD crossover loop: in the predicted structure, BubCD Y370 is situated alongside BubA’s extended C-terminus, with its hydroxyl group positioned to attack an adenylated BubA immediately upon formation (**Figure 4d-e**). This predicted structure may therefore represent a pre-reaction conformation of BubCD, while our crystal structures more likely represent an E1-E2 handoff state with BubCD Y370 juxtaposed to BubB S333.

We sought to reconstitute BubA-target conjugation *in vitro* using purified proteins. We first mixed purified *Citrobacter* BubA with BubCD in the absence and presence of ATP, and observed a high-molecular weight band that appeared specifically in the presence of ATP that may represent covalently-linked BubA and BubCD (**Figure 4f**, lanes 1-2). We next added BubB to the *in vitro* reactions, and observed a strong band at the combined molecular weight of BubA and BubB, again dependent on ATP (**Figure 4f**, lanes 3-4). This band was not present when we used a mutant BubB with its only lysine residue (K358) mutated to arginine (**Figure S4c**), suggesting that it represents a product with BubA covalently linked via its C-terminus to BubA K358. Mutation of the BubCD peptidase active site residue D104 did not affect catalysis, presumably because we used isolated BubA that does not require processing by BubCD (**Figure 4f**, lane 5). Mutation of BubCD R227 to alanine, which is predicted to disrupt BubA adenylation, reduced but did not eliminate the BubA-BubB product complex (**Figure 4f**, lane 6), while mutation of BubCD Y370 to phenylalanine eliminated the BubA-BubB product (**Figure 4f**, lane 7). Curiously, the high molecular weight BubA-BubCD species still formed with BubCD Y370F, suggesting that instead of representing an on-pathway BubA∼BubCD intermediate, it instead represents a minor off-pathway product. Finally, we found that a BubB S333A/H335A double mutant also eliminated the BubA-BubB product (**Figure 4f**, lane 8).

To complement *in vitro* experiments and test whether *Citrobacter* Type I Bub proteins can conjugate BubA to proteins in cells, we coexpressed His_6_-FLAG-tagged BubA, BubB, and BubCD in *E. coli* cells. We purified His_6_-FLAG-tagged BubA and any BubA-target conjugates in denaturing conditions using Ni^2+^ affinity, then analyzed by SDS-PAGE and western blotting against BubA (**Figure 4g**). When we coexpressed His_6_-FLAG-BubA, BubB, and wild-type BubCD, we observed a series of high-molecular weight bands likely corresponding to nonspecific BubA conjugation to *E. coli* proteins (**Figure 4g**, lane 3). Consistent with our *in vitro* data, mutation of BubCD adenylation active site residue R227 to alanine reduced these bands, and mutation of Y370 to phenylalanine eliminated them (**Figure 4g**, lanes 4-5). Similarly, mutation of BubB putative catalytic residues S333 and H335 to alanine also eliminated BubA-target conjugates (**Figure 4g**, lane 6).

### Type II Bub operons function through oxyester intermediates

Whereas Type I Bub operons encode a fused peptidase-E1 protein (BubCD), Type II Bub operons encode a predicted E2-E1 fusion protein (BubBD). We used AlphaFold 3 to predict the structures of Type II Bub operon proteins from *E. coli* ZDHYS365 (**Table S1**). We first predicted the structure of a BubBD dimer bound to two copies of BubA and two molecules of AMP, to mimic an adenylated BubA intermediate state. We found that the BubBD dimer is confidently predicted, and that BubA is predicted to bind the BubA E1 domain as in our structures of *Citrobacter* BubA bound to BubCD (**Figure S5a-c**). In these models, the position of the BubBD E2 domain is distinct from that observed in our structures of *Citrobacter* BubCD bound to BubB, with the predicted E2 active site positioned near the adenylation active site and crossover loop of the opposite BubBD protomer (**Figure S5c**). Like BubCD, BubBD proteins possess an invariant tyrosine in the crossover loop of their adenylation domains (Y342 in *E. coli* BubBD; **Figure S5g**). In a structure prediction including only the BubBD adenylation domain, BubA, and AMP, this tyrosine residue is predicted to be positioned close to the BubA C-terminus and AMP, similar to our predicted structure of the *Citrobacter* BubCD-BubA complex (**Figure S5d**). In a prediction of the full-length BubBD dimer bound to AMP (without BubA), BubBD Y342 is predicted to swing outward toward the predicted E2 active site of the opposite BubBD protomer (**Figure S5e**). Finally, in the prediction of a full-length BubBD dimer bound to BubA and AMP, BubBD Y342 is predicted to adopt a position outside both active sites (**Figure S5f**). These predictions suggest that the BubBD crossover loop is flexible, and that the putative catalytic tyrosine residue Y342 could access both the adenylation active site and the E2 active site.

To test whether Type II Bub operons mediate Ubl conjugation, we mixed *E. coli* BubA with BubBD *in vitro*. In the presence of ATP, we observed formation of high molecular-weight species likely representing BubA∼BubBD intermediates or off-pathway products, plus an apparent BubA-BubA covalent conjugate (**Figure S5h**). Disruption of BubBD adenylation activity (R219A) or mutation of the protein’s predicted E1 or E2 catalytic residues (Y342F or S106A, respectively) eliminated formation of the putative BubA-BubA conjugate (**Figure S5h**). Overall, these data support the idea that both Type I and Type II Bub operons function via oxyester intermediates, using a conserved tyrosine residue on their adenylation domain crossover loops, and a conserved serine in their E2 domains.

### BubE is a serine protease

All Bub operons possess *bubE*, which encodes an uncharacterized protein with a DUF6527/PF20137 domain. We were unable to determine a structure of BubE from any Bub operon, so we instead performed AlphaFold 3 structure predictions of *Citrobacter* and *E. coli* ZDHYS365 BubE proteins (**Figure S6a-d**). The structure predictions reveal BubE as a compact β-sandwich domain protein with a flexible N- or C-terminal extension containing a high fraction of solvent-exposed hydrophobic residues (**Figure 5a, Figure S6a-b**). These extensions are not predicted to form transmembrane α-helices, but they may mediate peripheral association of BubE with membranes. A cluster of four conserved cysteine residues (C57/C59/C61/C99 in *Citrobacter* BubE) suggest that the protein binds a structural zinc ion (**Figure 5a-c**). Neither the DALI^24^ nor Foldseek^25^ structure-comparison servers identified close structural homologs of BubE.

**Figure 5.**
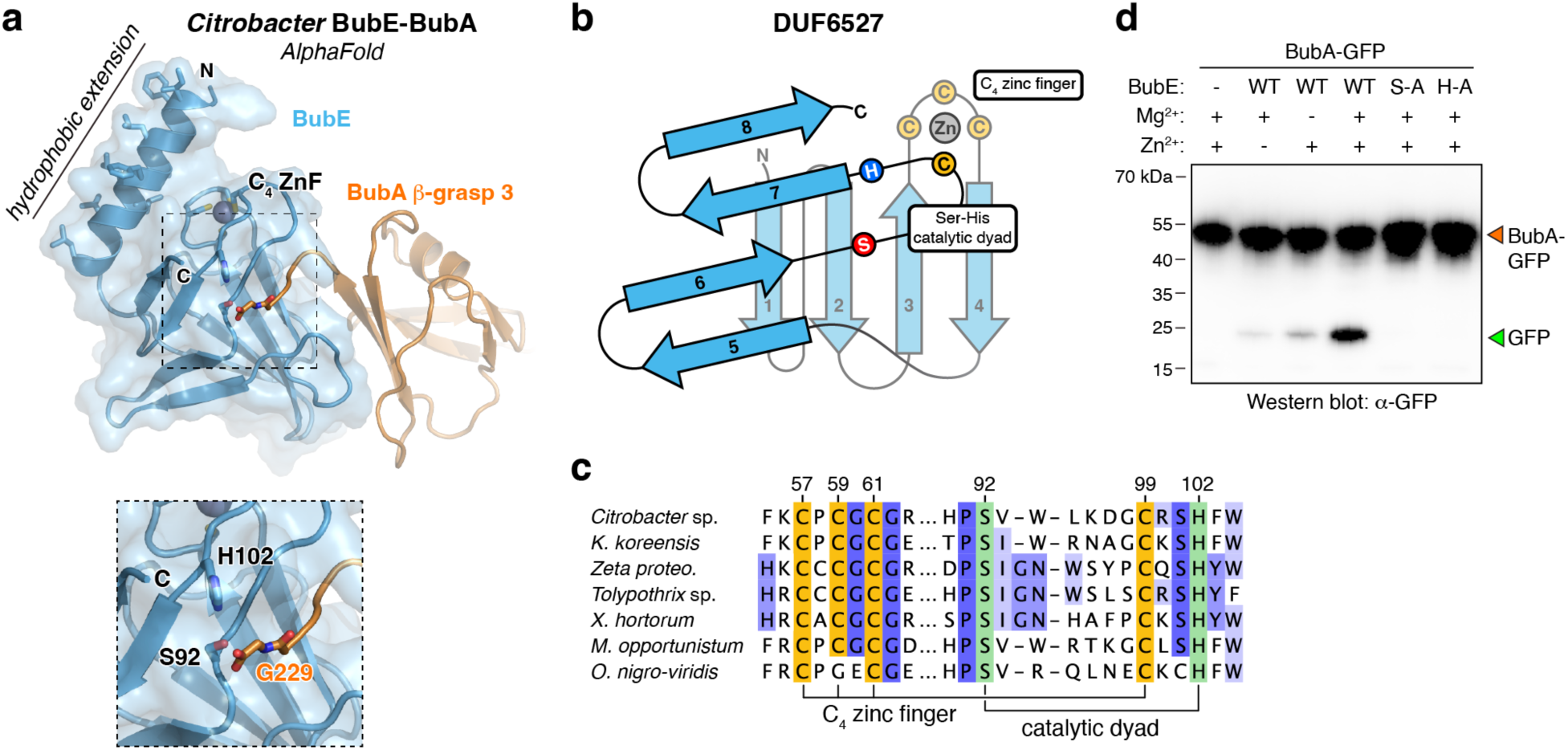
BubE is a serine protease. **(a)** AlphaFold 3 predicted structure of *Citrobacter* BubE (blue) bound to BubA (orange; only the third β-grasp domain of BubA shown) and Zn^2+^ (gray sphere). The N-terminal hydrophobic extension of BubE is labeled with hydrophobic residues shown as sticks, and the putative active site is indicated with a dotted box. *Inset bottom:* Closeup view showing that the C-terminal residue of BubA (G229) is predicted to be positioned close to BubE S92 and H102. (b) Schematic of the DUF6527 fold, with β-strands numbered 1-8, cysteine residues involved in the C_4_ zinc finger shown in yellow, and the serine/histidine catalytic dyad shown in red and blue. **(c)** Section of sequence alignment of 544 BubE (DUF6527) proteins from Type I and Type II Bub operons. Highlighted in green are the invariant serine (S92 in *Citrobacter* BubE) and histidine (H102) proposed to make up the catalytic dyad, and shown in yellow are the cysteine residues (C57, C59, C61, and C99) involved in the C_4_ zinc finger. BubE proteins shown are from *Citrobacter* sp. RHBSTW-00271 (IMG 2938140954), *Kangiella koreensis* SW-125 (IMG 644987649), *Zeta proteobacterium* SCGC AB-604-B04 (IMG 2264879631), *Tolypothrix* sp. PCC 7601 (IMG 2501546804), *Xanthomonas hortorum* pv. gardneri 7 (IMG 2501866951), *Mesorhizobium opportunistum* WSM2075 (IMG 2503204319), and *Oscillatoria nigro-viridis* PCC 7112 (IMG 2504086280) (see **Table S1**). **(d)** *In vitro* cleavage of BubA-GFP fusion by *Citrobacter* BubE wild-type (WT), S92A (S-A), and H102A (H-A) in the presence or absence of Zn^2+^ (10 µM ZnCl_2_) and Mg^2+^ (20 mM MgCl_2_). Proteins are detected by anti-GFP western blot.

AlphaFold 3 confidently predicted that both *Citrobacter* and *E. coli* BubE bind their cognate BubA proteins (**Figure S6a-d**). In both cases, BubE is predicted to bind the C-terminal β-grasp domain of BubA, with the conserved C-terminal residue of BubA (G229 in *Citrobacter* BubA) positioned close to a pair of invariant residues (S92 and H102 in *Citrobacter* BubE; **Figure 5a-b**). The presence of this conserved residue pair in BubE, and their predicted position close to the BubA C-terminus, suggests that BubE functions either in Ubl transfer (as in E2 enzymes) or as a serine protease. Typical serine proteases possess a catalytic triad comprising serine, histidine, and aspartate residues, but some serine proteases lack the aspartate residue and perform proteolysis with a serine-histidine catalytic dyad^26^.

We tested whether *Citrobacter* BubE could act as a BubA-specific peptidase by incubating purified *Citrobacter* BubE with a BubA-GFP fusion protein. In the presence of wild-type BubE, but not S92A or H102A mutants, we detected a specific cleavage product indicative of proteolysis of BubA-GFP (**Figure 5d**). Extended incubations (48 hours at room temperature) were required to observe BubA-GFP cleavage, showing that BubE’s catalytic activity is extremely weak, at least *in vitro*. We found that BubE is most active in a buffer containing Zn^2+^, in keeping with its predicted C_4_ zinc finger motif (**Figure 5d**), and at near-neutral pH (**Figure S6d**). We hypothesize that the low *in vitro* catalytic activity of *Citrobacter* BubE may be related to the hydrophobicity of its N-terminal extension and resultant tendency to aggregate in solution; attempts to truncate BubE to remove this extension yielded insoluble protein, preventing us from testing this idea. Nonetheless, our structure predictions and biochemical data show that BubE has peptidase activity attributable to its conserved serine-histidine catalytic dyad, thus revealing the DUF6527 domain to be a novel family of serine protease.

To determine BubE’s role in BubA-target conjugation, we expressed BubE with His_6_-FLAG-tagged BubA, BubB, and BubCD in *E. coli* cells. Expression of wild-type BubE almost completely eliminated the observed BubA-target conjugates in this experiment (**Figure 4g**, lane 7), consistent with a role for BubE in cleaving BubA-target isopeptide-linked conjugates to negatively regulate the Bub operon. Consistent with our *in vitro* cleavage data, mutation of S92 or H102 to alanine resulted in reappearance of BubA-target conjugates (**Figure 4g**, lanes 8-9).

## Discussion

Bacteria possess a variety of protein conjugation pathways related to eukaryotic ubiquitination pathways, the biochemical mechanisms and biological roles of which are only beginning to be understood. Type II CBASS pathways, along with related Type II Pycsar and MBL pathways, use a fused E2-E1 protein that is related to eukaryotic Atg7 to conjugate their respective signaling enzymes to unknown targets to activate antiphage immunity^2,3,27^. Type I and II Bil pathways, meanwhile, use separate eukaryotic-like E1 and E2 proteins to conjugate a Ubl to phage proteins and thereby interrupt virion assembly and infectivity^4,5,15^. While bacterial Type II CBASS/Pycsar/MBL pathways and Bil pathways are clearly evolutionarily linked to eukaryotic noncanonical and canonical ubiquitination pathways, respectively, here we outline the architecture and biochemistry of a more diverged set of bacterial pathways that we term Bub (<u>B</u>acterial <u>ub</u>iquitination-like).

Our structural and biochemical data show that Bub operons encode Ubl, E1, E2, and at least one peptidase protein, which together constitute a *bona fide* protein conjugation pathway. The most striking feature of Bub operons is that they catalyze protein conjugation without catalytic cysteine residues. Distinct from all other ubiquitination-related pathways, Bub operons utilize Ubl∼tyrosine and Ubl∼serine oxyester intermediates instead of Ubl∼cysteine thioesters. Oxyesters are typically much less reactive than thioesters, to the extent that Ubl∼E2 covalent complexes with E2 catalytic cysteine-to-serine mutations have been trapped and visualized by X-ray crystallography^28–30^. Nonetheless, serine-linked Ubl∼E2 oxyesters have been shown to retain sufficient reactivity for Ubl transfer to E3 proteins^29^. In Bub E1 and E2 proteins, the reactivity of the oxyester intermediates is likely supported by structural features of their active sites, including the conserved accompanying residues (histidine+aspartate in Type I Bub; lysine+histidine in Type II Bub) near the E2 catalytic serine.

All characterized bacterial ubiquitination-related pathways including Type II CBASS/Pycsar/MBL and Type I/II Bil encode a JAB/JAMM family peptidase, which negatively regulates signaling by cleaving substrate-target conjugates^2,4^ and/or cleave pre-proteins to expose reactive C-termini for adenylation and eventual target conjugation^4^. Type I Bub operons encode a JAB/JAMM family peptidase at the N-terminus of their BubCD proteins, and we show that this domain can cleave the BubAB pre-protein. Notably, both our *in vitro* and *in vivo* ubiquitination assays with *Citrobacter* Bub proteins used wild-type BubCD proteins with an intact peptidase active site, and we nonetheless observed robust BubA-target conjugation in both cases (**Figure 4f, i**). These data suggest that the JAB/JAMM peptidase domain of BubCD in Type I Bub operons may function primarily or solely to cleave the BubAB pre-protein, rather than negatively regulate the pathway by cleaving BubA-target conjugates.

While only Type I Bub operons encode a JAB/JAMM family peptidase, all Bub operons encode BubE. We show that BubE is a novel serine protease with a conserved serine-histidine catalytic dyad. Serine proteases constitute the first and best-documented case of convergent evolution, with at least 15 unrelated protein scaffolds independently evolving this enzymatic activity^31^.

While typical serine proteases use a catalytic triad comprising serine, histidine, and aspartate residues, some families lack the aspartate and possess either a serine-histidine or serine-lysine catalytic dyad^26^. Our data show that BubE is an active peptidase and suggest that it negatively regulates Bub operon activity by cleaving BubA-target conjugates, similar to how the Cap3 peptidase negatively regulates Type II CBASS signaling by cleaving conjugates produced by the operon’s E2-E1 ligase protein Cap2 (ref. ^2^). The presence of N- or C-terminal extensions with a high fraction of hydrophobic residues, which are conserved across BubE proteins, further suggests that BubE could associate with membranes to spatially regulate its activity.

Thus far, we have been unable to identify a Bub operon that confers protection against bacteriophage infection in *E. coli* (data not shown), leaving open the question of these pathways’ biological role(s). Our analysis of Bub operons’ genomic context indicates that they tend to be found in mobile elements and/or defense loci: for example, both the *Citrobacter* and *E. coli* Bub operons under study here are located in mobile elements inserted at tRNA genes in their host genomes (**Figure S7**). We also find that a large fraction of Bub operons are associated with the DNA damage-sensitive transcription factors CapW/BrxR or CapH+CapP (**Table S1**), both of which are also associated with a variety of defense pathways like CBASS, DISARM, and Wadjet^17,18,32,33^. Together, these data suggest that Bub operons play roles in antiphage immunity or stress response. Future work will be needed to define the specific targets of Bub-mediated Ubl conjugation, and the biological role(s) of this modification in bacterial cells.

## Materials and Methods

### Protein expression and purification

All proteins used in this study are listed in **Table S2**. For each protein, codon-optimized genes were individually cloned into *E. coli* expression vectors encoding an N-terminal TEV protease-cleavable His_6_-tag (UC Berkeley Macrolab vector 2B-T, Addgene ID 29666) or His_6_-MBP-tag (UC Berkeley Macrolab vector 2C-T, Addgene ID 29706). For coexpression, multigene coexpression cassettes were generated by PCR and cloned into UC Berkeley Macrolab vector 2B-T to generate an N-terminal TEV protease-cleavable His_6_-tag on BubA; PCR mutagenesis was used to generate point mutations and to insert the FLAG sequence (DYKDDDDK) into the His_6_-BubA expression vector for western blotting.

Vectors were transformed into *E. coli* Rosetta 2 pLysS (EMD Millipore), and 1L cultures were grown at 37°C in 2XYT media (RPI) to an OD_600_ of 0.6 before induction with 0.25 mM IPTG at 20°C for 16-18 hours. Cells were harvested by centrifugation, resuspended in buffer A (25 mM Tris-HCl pH 8.5, 300 mM NaCl, 5 mM MgCl_2_, 10% glycerol, and 5 mM β-mercaptoethanol) containing 5 mM imidazole, then lysed by sonication (Branson Sonifier). Lysates were clarified by centrifugation, then supernatants were passed over a Ni-NTA Superflow column (Qiagen) in resuspension buffer. The column was washed in buffer A containing 20 mM imidazole, then eluted in buffer A containing 400 mM imidazole. Eluates were concentrated and buffer-exchanged by ultrafiltration (Amicon Ultra; EMD Millipore). His_6_-tagged proteins were cleaved with TEV protease (expressed and purified in-house from plasmid pRK793, Addgene ID 8827)^34^ then passed over a HisTrap HP column (Cytiva) to remove His_6_-tagged TEV protease and uncleaved proteins. Untagged proteins were further passed over a Superdex 200 Increase size exclusion column (Cytiva) in size exclusion buffer (25 mM Tris-HCl pH 8.5, 300 mM NaCl, 5 mM MgCl_2_, 10% glycerol, and 1 mM DTT). For His_6_-MBP-tagged *Citrobacter* BubE, samples were injected into a Hitrap Q HP column (Cytiva) and eluted with a 0.1-1 M NaCl gradient, then passed over a Superose 6 Increase size exclusion column (Cytiva). Peak fractions were concentrated and buffer exchanged by ultrafiltration and stored at 4°C for crystallography or - 80°C for biochemical assays.

For denatured purification of His_6_-FLAG-tagged *Citrobacter* BubA after coexpression with other Bub proteins, proteins were purified by Ni-NTA affinity chromatography as above in buffers containing an additional 8M urea.

### Ubiquitination assays

For *in vitro* assays, all proteins were purified individually, then 10 µg of each protein was mixed in 20 µL reaction buffer (10 mM Tris-HCl pH 8.5, 200 mM NaCl, 20 mM MgCl_2_, and 1 mM TCEP (tris(2-carboxyethyl)phosphine), plus 2 mM ATP as indicated. Samples were incubated at 37°C for 30 minutes, then analyzed by SDS-PAGE with Coomassie blue staining or western blotting.

For *in vivo* assays, *Citrobacter* His_6_-Flag-BubA, BubB (wild type or mutants) and BubCD (wild type or mutants) were cloned into a single coexpression vector, and BubE (wild type or mutants) was cloned separately. Proteins were transformed into *E. coli* Rosetta2 pLysS cells and expressed as described above, then single-step Ni-NTA purification was performed in denaturing conditions and samples analyzed by SDS-PAGE with Coomassie blue staining and western blotting.

### BubC in vitro activity assays and N-terminal sequencing

To the peptidase activity of BubCD, purified *Citrobacter* BubAB (10 µg) was mixed with 10 µg of full-length BubCD or BubCD 2-164 (wild type or mutants) in 20 µL reaction buffer (10 mM HEPES pH 7.5, 100 mM NaCl, 20 mM MgCl_2_, 20 µM ZnCl_2_, and 1 mM TCEP), then incubated at 37°C for 30 minutes. Reactions were analyzed by SDS-PAGE with Coomassie blue staining.

For protein N-terminal sequencing by Edman degradation, cleavage products were separated by SDS-PAGE, transferred to a Bio-Rad ImmunBlot PVDF membrane (Bio-Rad Turbo Transfer Kit, 170-4272), and visualized by staining with Coomassie Blue R-250. The band representing the C-terminal cleavage product of BubAB was excised, the membrane was washed with methanol and deionized water, then loaded onto a Shimadzu PPSQ-53A instrument (Iowa State University Protein Facility) for analysis. Five rounds of Edman degradation were performed, and the “evaluated value” scores for each round were analyzed to obtain the likely N-terminal sequence (**Figure S1**).

### BubE in vitro activity assays

To generate a model substrate for BubE activity assays, BubA was cloned into a vector encoding a C-terminal GFP tag (UC Berkeley Macrolab vector H6-msfGFP, Addgene ID 29725) and purified as above. The BubA-GFP fusion protein (10 µg) was mixed with 10 µg of BubE (wild type or mutants) in 20 µL reaction buffer (10 mM HEPES pH 7.5, 100 mM NaCl, 20 mM MgCl_2_, 20 µM ZnCl_2_, and 1 mM TCEP), then incubated at room temperature for 48 hours. Reactions were analyzed by western blotting.

### Western blots

For western blotting, proteins were separated on 4-20% Mini-PROTEAN TGX gels (BIO-RAD 4561096), then transferred to PVDF membranes (BIO-RAD Turbo Transfer Kit, 170-4272). The membranes were blocked with 10 mL 5% non-fat dry milk in TBST (50 mM Tris-HCl pH 7.5, 150 mM NaCl, 0.1% Tween-20) with gentle shaking at room temperature for 1 hour, following by incubation with a 1:3,000 dilution primary antibody (anti-GFP from mouse lgG; Roche 11814460001) with 10 mL 5% milk in TBST overnight at 4°C or 1 hour at room temperature.

After washing three times with TBST, the blots were incubated with 10 mL HRP-linked mouse secondary antibody (Millipore Sigma 12-349) at a 1:4,000 dilution in TBST at room temperature for 1 hour. The membranes were washed three times with TBST, incubated in ECL detection reagent (Cytiva RPN2232) for 1 minute, and then imaged on a Bio-Rad ChemiDoc imager. For the Flag-tagged samples, the same western blot procedure was followed, except the primary antibody was changed to a 1:16,000 dilution anti-Flag mouse lgG (Millipore Sigma, F3165).

### Crystallography

For crystallization of *Citrobacter* BubCD, purified protein at 16 mg/mL in crystallization buffer (20 mM HEPES pH 7.5, 200 mM NaCl, 5 mM MgCl_2_, and 1 mM TCEP) was mixed 1:1 with well solution containing 100 mM HEPES pH 7.5, 0.2 M K/Na tartrate, 10% Isopropanol, 20% PEG 3350 in a hanging drop format. Crystals were harvested into a cryoprotectant solution containing an additional 10% glycerol and frozen in liquid nitrogen. The 1.8 Å resolution dataset was collected at the SSRL beamline 9-2 on August 16, 2023 (**Table S3**). The dataset was processed with the autoxds pipeline, which uses XDS^35^ for integration, AIMLESS^36^ for scaling, and TRUNCATE^37^ for conversion to structure factors. The structure was determined by molecular replacement in PHASER^38^ using a predicted structure from AlphaFold 2 (ref. ^22^). The model was manually rebuilt in COOT^39^ and refined in phenix.refine^40^ using positional and individual isotropic B-factor and domain-based anisotropic TLS refinement.

For crystallization of *Citrobacter* BubCD^D104A^, purified protein at 30 mg/mL in crystallization buffer was mixed 1:1 with well solution containing 100 mM Tris pH 8.5, 0.1 M NaOAc, 10% Isopropanol, 28% PEG 3350 in a hanging drop format. Crystals were harvested into a cryoprotectant solution containing an additional 10% glycerol and frozen in liquid nitrogen. The 2.0 Å dataset was collected at the SSRL beamline 9-2 on August 16, 2023 (**Table S3**). The dataset was processed with the autoxds pipeline, and the structure was determined by molecular replacement in PHASER using the wild type BubCD structure. The model was manually rebuilt in COOT and refined in phenix.refine using positional and individual isotropic B-factor and domain-based anisotropic TLS refinement.

For crystallization of the *Citrobacter* BubCD^D104A^-BubA^155–229^, the two purified proteins in crystallization buffer were mixed in a 1:1.2 ratio with 2 mM AMPCPP (Millipore Sigma, M6517), incubated on ice for 10 minutes, and then mixed 1:1 with well solution containing 100 mM MES pH 6.0, 2.2 M NaOAc pH 7.0 in a hanging drop format. Crystals were harvested into a cryoprotectant solution containing an additional 30% glycerol and frozen in liquid nitrogen. The 1.95 Å dataset was collected at the SSRL beamline 12-2 on April 13, 2024 (**Table S3**). The dataset was processed with the autoxds pipeline, and the structure was determined by molecular replacement in PHASER using the BubCD^D104A^ structure we solved and a predicted Ubl structure from AlphaFold2. The model was manually rebuilt in COOT and refined in phenix.refine using positional and individual isotropic B-factor and domain-based anisotropic TLS refinement.

For crystallization of the *Citrobacter* BubCD^D104A^-BubB, the two purified proteins in crystallization buffer were mixed in a 1:1.2 ratio, incubated on ice for 10 minutes, and then added an equal volume of well solution containing 100 mM sodium cacodylate pH 7.0, 0.2 M MgCl_2_, 10% PEG 3000 in a sitting drop format. Crystals were harvested into a cryoprotectant solution containing an additional 20% glycerol and frozen in liquid nitrogen. The 2.53 Å dataset was collected at the SSRL beamline 12-1 on July 10, 2024 (**Table S3**). The dataset was processed with the autoxds pipeline, and the structure was determined by molecular replacement in PHASER using the BubCD^D104A^ structure we solved and a predicted E2 structure from AlphaFold2. The model was manually rebuilt in COOT and refined in phenix.refine using positional and individual isotropic B-factor and domain-based anisotropic TLS refinement.

For crystallization of *Citrobacter* BubCD^D104A/Y370F^-BubB-BubA^155–229^, the three purified proteins in crystallization buffer were mixed in a 1:1.1:1.2 ratio with 2 mM ATP, incubated on ice for 10 minutes, and then mixed 1:1 with well solution containing 0.24 M sodium malonate pH 7.0 and 15% PEG 3350 in hanging drop format. Crystals were harvested into a cryoprotectant solution containing an additional 15% glycerol and frozen in liquid nitrogen. A 1.93 Å resolution dataset was collected at the SSRL beamline 12-1 on May 29, 2024 (**Table S3**). The dataset was processed with the autoxds pipeline, and the structure was determined by molecular replacement in PHASER by separately placing our structures of BubCD^D104A^, BubA^155–229^, and BubB. In refined electron density maps produced after placing two BubCD protomers, one BubA protomer, and one BubB protomer, we noticed strong difference density corresponding to a second BubB protomer in a position that would overlap itself due to symmetry. We manually placed this protomer based on its orientation with respect to BubCD, then refined it in phenix.refine with an occupancy of 0.5 to account for the fact that each asymmetric unit has this protomer oriented in one of two possible ways (using the phenix.refine argument “allow_polymer_cross_special_position=True”). To minimize non-ideal geometry in the affected protomer, refinement was performed in two stages. First, positional and isotropic B-factor refinement was performed on the full model, yielding *R* and *R_free_* values of 17.86% and 20.17%, but with the resulting model showing significant geometry outliers in the affected BubB protomer. After that, the affected BubB protomer model was replaced with a copy of the unaffected BubB protomer, its occupancy set to 0.5, then a final round of refinement was performed with rigid-body refinement of the affected BubB protomer only, with isotropic B-factor and TLS refinement for the full model. This yielded both better final *R* and *R_free_* values (17.60% and 19.64%, respectively) and significantly better overall geometry.

## Acknowledgements

The authors thank members of the Corbett lab and A. Whiteley for helpful discussions and critical reading of the manuscript. The authors acknowledge funding from the National Institutes of Health (R35 GM144121 to K.D.C.) and from the Howard Hughes Medical Institutes Emerging Pathogens Initiative (to K.D.C.). L.R.C. is supported by the UCSD Molecular Biophysics Training Grant (T32 GM139795). Use of the Stanford Synchrotron Radiation Lightsource, SLAC National Accelerator Laboratory, is supported by the U.S. Department of Energy, Office of Science, Office of Basic Energy Sciences under Contract No. DE-AC02-76SF00515. The SSRL Structural Molecular Biology Program is supported by the DOE Office of Biological and Environmental Research, and by the National Institutes of Health, National Institute of General Medical Sciences (including P41GM103393).

