## Supplemental Information for "Mechanistic basis for protein conjugation in a diverged bacterial ubiquitination pathway"

#### Table S1. Bub operons

*See separate Excel workbook*

### Table S2. Proteins used in this study

| **Species** | **Gene** | **Accession** | **Sequence** |
|---|---|---|---|
| *Citrobacter* sp. RHBSTW-00271 | *bubAB* | IMG 2938140956 (bubA in black; bubB in red) | MQDIQSQHHHRRFIEVADETLSFRQVVMEDSTPNGSQISAASGFKPDQM<br>PVVLMLLPNGSLEDIRPDEVVDLSSEVRRFIVVESDRTYFFTIDGARLEWP<br>CRFITGYSIRQLGDIGDNKKLLLEREDEADLEVQNDQIIDLDGDGIERFISR<br>KATWKLNIQGKEFTFDTPTVVIRDAVIRAGLNPNQAWHIFLKVEGQPKVEK<br>NIDDVIDLRTPGIEKLRLTPKDVNNGEEPRATRRDFSLRPEDEHYLDEMGY<br>CWETRLVGNARWLIIHDYELPDGYNHHQVNLALLITSGYPVNMLDMFYVY<br>PPLVRVNGVNIPATEATVAIDSVAYQRWSRHRSWNPEIDSVISQLAMADG<br>CLQKEVGQ |
| *Citrobacter* sp. RHBSTW-00271 | *bubCD* | IMG 2938140955 (bubC in black; bubD in red) | MSSGNRLILTQELHTMLQKHLFPGDGKEAAAILICNRYEGGRLKLLAKELIL<br>VPYEECKSRTSDFIAWPGNYLEKAIDVAEEKSMSIILIHSHPGGFLVFSDTD<br>DSSDMQTMQSLFQGVDAIHGSAIMIHSGEMRARLYREGKFAENVELVTV<br>AGDDIHYWWDDKTEQQLKPIAFTSGMTDTFQKLTAAIIGVSGTGSIVAEQV<br>ARLGFGEILLIDHDHIEKKNLNRILNSTLKDALSHRPKVDMFAEAIRCIRGED<br>ISRPINNTIFSREAVLAAANADVLFCCVDTYLARMIADRIASSFLIPLLDVGV<br>KIPTHVDPDDGRKITDVTGRIDYVKPGGSTLSDRLVYTPELIYRENLNAEE<br>YEEQLERGYITGVEEEAPSVITLNMRAASACVSEFIARCFPFREYPNKRFT<br>RTFFSLAGVEEDYIDESSITQALNTRLAVGGEEPLLGLPELGDK |
| *Citrobacter* sp. RHBSTW-00271 | *bubE* | IMG 2938140954 | MYIFSKLFYFLLRKLEHLLPARRLVIINQDSLPEKMPLRSIILARDDDEDWCI<br>GFKCPCGCGRTVELLVIDEASPRWDYCLDTNSLPSLHPSVWLKDGCRSH<br>FWIKKGRVFWV |
| *E. coli* ZDHYS365 | *bubA* | NCBI WP_053069300.1 | MKNKFIKINDKIVEIDDLTPTGAQILLVAGVKDIVEYVLFQKLKNGLLEEIRPE<br>EKTTLDKEGVETFLMFNSDRTYRFTLNGKMFDWGAPSLTGATVKSLAEC<br>DFDSNDVWLEMKNEKDKLITDHEHIDLTQPNVERLYTQETSINIIVNAKMR<br>VVNRRVISYWDVVHLAYEHAENKETSIYSVDYAKGPISNPEGSMVDGQYV<br>QLTDGMIFYVTQTDKS |
| *E. coli* ZDHYS365 | *bubBD* | NCBI WP_063617865.1 (bubB in black; bubD in red) | MSLKLINLNPCLKRLQDEGYELEIRDGFLMVHSVPYVNSNKEVLKGTLITN<br>LVLVSPDVVGKPNTHQMYFNGEHPCHPDGRILTAIQHTTPNQTLGNGIIG<br>NHWFSNKPPSGSYDDYYHQVTSYVRVIESQAKAIDENVSAKTFISHYLTN<br>DEDIFTYQDTASARIGTVQLNNLIANQKLAIIGLGGTGSYILDMISKTCVSEI<br>HLYDKDQFQQHNSFRAPGAASIDDLEKKQTKCDYYYQTYSNLRNGVIPH<br>NEMLSETNLHELINYDFIFISIDSATSRKMITDYLYQHKISFIDVGMGLSFTE<br>DKSSIFGTCRSTLYTKDMDDTPLKSLPTISRDDEVYRSNIQLVELNCLNAA<br>FAVIQWKKLFGFYCDDMSSYEMTYSLGLNKIANKVMERVE |
| *E. coli* ZDHYS365 | *bubE* | NCBI WP_074146335.1 | MKTLSITFTEFIPEQLEEGILYVSMPYSTIAHLCACGCKNEVITPLSPLDWS<br>MTYNGKEISLHPSIGNWQFPCRSHYWIRGNKVVWAGNMHNDIIENNRSA<br>NLALKMKKSETNITSHNYTETGMNDKPKQTSYINQLMNTIREWLGYK |

**Table S3 (part 1). Crystallographic data collection and structure determination**

| | BubCD wild-type | BubCD$^{D104A}$ | BubCD$^{D104A}$-BubA$^{155-229}$ |
|---|---|---|---|
| **Data collection** | | | |
| Data collection date | August 16, 2023 | August 16, 2023 | April 13, 2024 |
| Beamline | SSRL 9-2 | SSRL 9-2 | SSRL 12-2 |
| Wavelength (Å) | 0.97946 | 0.97946 | 0.97946 |
| Space group | $P3_121$ | $P2_1$ | $P4_32_12$ |
| Cell dimensions | | | |
| *a*, *b*, *c* (Å) | 78.50, 78.50, 159.29 | 65.87, 73.06, 98.40 | 96.64, 96.64, 135.20 |
| α, β, γ (°) | 90, 90, 120 | 90, 108.72, 90 | 90, 90, 90 |
| Resolution (Å)* | 39.25-1.80 (1.84-1.80) | 39.29-2.00 (2.05-2.00) | 39.31-2.34 (2.42-2.34) |
| $R_{merge*}$ | 0.118 (1.423) | 0.123 (0.893) | 0.060 (1.634) |
| $I / \sigma I$ | 14.3 (1.9) | 12.0 (2.2) | 25.8 (1.7) |
| Completeness (%) | 99.9 (99.5) | 98.9 (98.2) | 100 (100) |
| Redundancy | 10.0 (9.0) | 6.9 (6.8) | 13.4 (12.7) |
| **Refinement** | | | |
| Resolution (Å) | 39.25 - 1.80 | 39.29 - 2.00 | 39.31 - 2.34 |
| No. reflections | 53435 | 59117 | 27686 |
| $R_{work}$ (%) | 15.88 (21.35) | 17.22 (22.66) | 24.19 (31.67) |
| $R_{free}$ (%) | 18.49 (25.27) | 20.71 (25.94) | 26.89 (35.10) |
| No. atoms | | | |
| Protein (non-H) | 3468 | 6762 | 4137 |
| Ligand/ion (non-H) | 2 ($Na^+$, $Zn^{2+}$) | 14 ($Na^+$, acetate) | 0 |
| Water | 533 | 673 | 80 |
| Hydrogen | 3434 | 6754 | 4135 |
| *B*-factors | | | |
| Protein | 29.36 | 32.16 | 86.57 |
| Ligand/ion | 29.17 | 35.14 | - |
| Water | 38.87 | 34.59 | 74.33 |
| R.m.s. deviations | | | |
| Bond lengths (Å) | 0.007 | 0.003 | 0.005 |
| Bond angles (°) | 0.847 | 0.573 | 0.678 |
| Validation | | | |
| MolProbity score | 1.18 | 1.02 | 1.68 |
| Clashscore | 2.61 | 1.18 | 8.10 |
| Poor rotamers (%) | 0.53 | 0.27 | 0.89 |
| Ramachandran plot | | | |
| Favored (%) | 97.29 | 96.87 | 96.36 |
| Allowed (%) | 2.71 | 2.90 | 3.64 |
| Disallowed (%) | 0 | 0.23 | 0 |
| PDB ID | 9EA4 | 9EA5 | 9EA8 |
| SBGrid Data Bank ID | 1141 | 1142 | 1143 |

*Values in parentheses are for highest-resolution shell.

**Table S3 (part 2). Crystallographic data collection and structure determination**

| | BubCD$^{D104A}$-BubB | BubCD$^{D104A/Y370F}$-BubB-BubA$^{155-229}$ |
|---|---|---|
| **Data collection** | | |
| Data collection date | July 10, 2024 | May 29, 2024 |
| Beamline | SSRL 12-1 | SSRL 12-1 |
| Wavelength (Å) | 0.97946 | 0.97946 |
| Space group | $P4_32_12$ | $P4_32_12$ |
| Cell dimensions | | |
| *a*, *b*, *c* (Å) | 135.35, 135.35, 172.78 | 136.84, 136.84, 172.82 |
| α, β, γ (°) | 90, 90, 90 | 90, 90, 90 |
| Resolution (Å) | 106.55-2.53 (2.57-2.53) | 39.45-1.93 (1.96-1.93) |
| $R_{merge}$ | 0.110 | 0.080 |
| $I / \sigma I$ | 14.2 (1.6) | 17.3 (1.6) |
| Completeness (%) | 100 (100) | 100 (100) |
| Redundancy | 13.6 (13.9) | 13.8 (13.2) |
| **Refinement** | | |
| Resolution (Å) | 49.35-2.53 | 34.21-1.93 |
| No. reflections | 54374 | 122974 |
| $R_{work}$(%) | 24.64 (41.61) | 18.31 (29.59) |
| $R_{free}$(%) | 27.11 (43.33) | 20.36 (31.47) |
| No. atoms | | |
| Protein (non-H) | 8108 | 9680 |
| Ligand/ion (non-H) | 2 ($Na^+$) | 25 ($Na^+$, AMP) |
| Water | 156 | 646 |
| Hydrogen | 8004 | 0 |
| *B*-factors | | |
| Protein | 83.34 | 53.28 |
| Ligand/ion | 83.89 | 39.65 |
| Water | 77.07 | 49.08 |
| R.m.s. deviations | | |
| Bond lengths (Å) | 0.002 | 0.015 |
| Bond angles (°) | 0.521 | 1.232 |
| Validation | | |
| MolProbity score | 1.23 | 1.18 |
| Clashscore | 1.86 | 2.34 |
| Poor rotamers (%) | 1.14 | 0.38 |
| Ramachandran plot | | |
| Favored (%) | 96.40 | 97.04 |
| Allowed (%) | 3.50 | 2.88 |
| Disallowed (%) | 0.10 | 0.08 |
| PDB ID | 9EAE | 9EAV |
| SBGrid Data Bank ID | 1144 | 1145 |

*Values in parentheses are for highest-resolution shell.

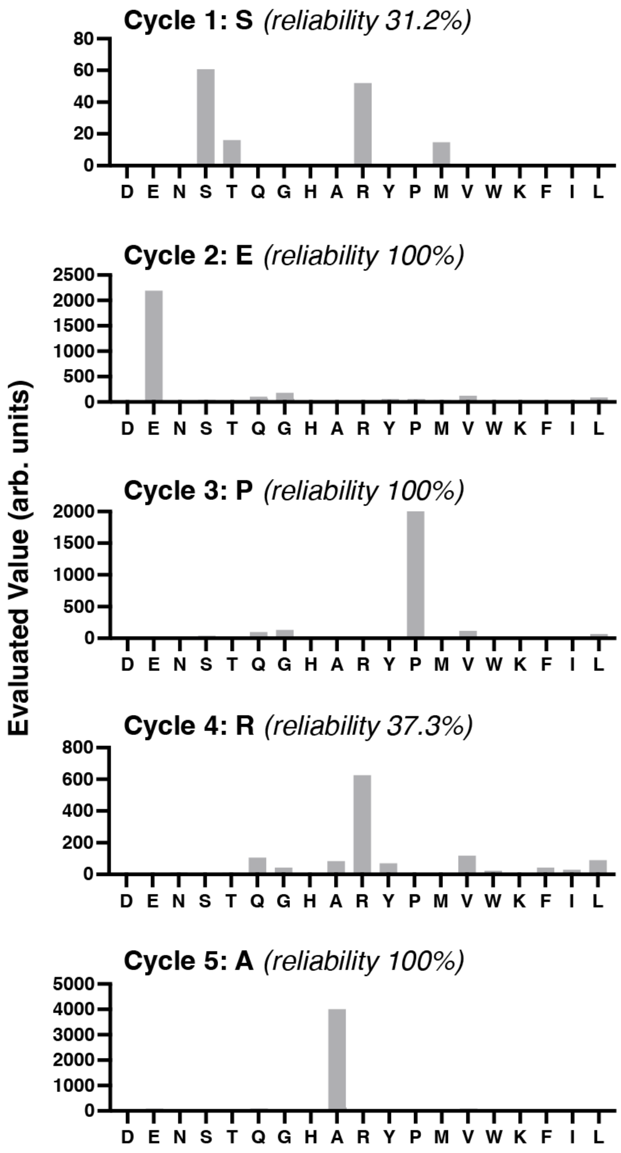

**Figure S1. Edman degradation analysis of BubAB cleavage product**

Graphs show amino acids (indicated by one-letter codes) detected in cycles 1-5 of Edman degradation of the C-terminal cleavage product of BubCD-mediated cleavage of BubAB (dotted box in **Figure 1b**)

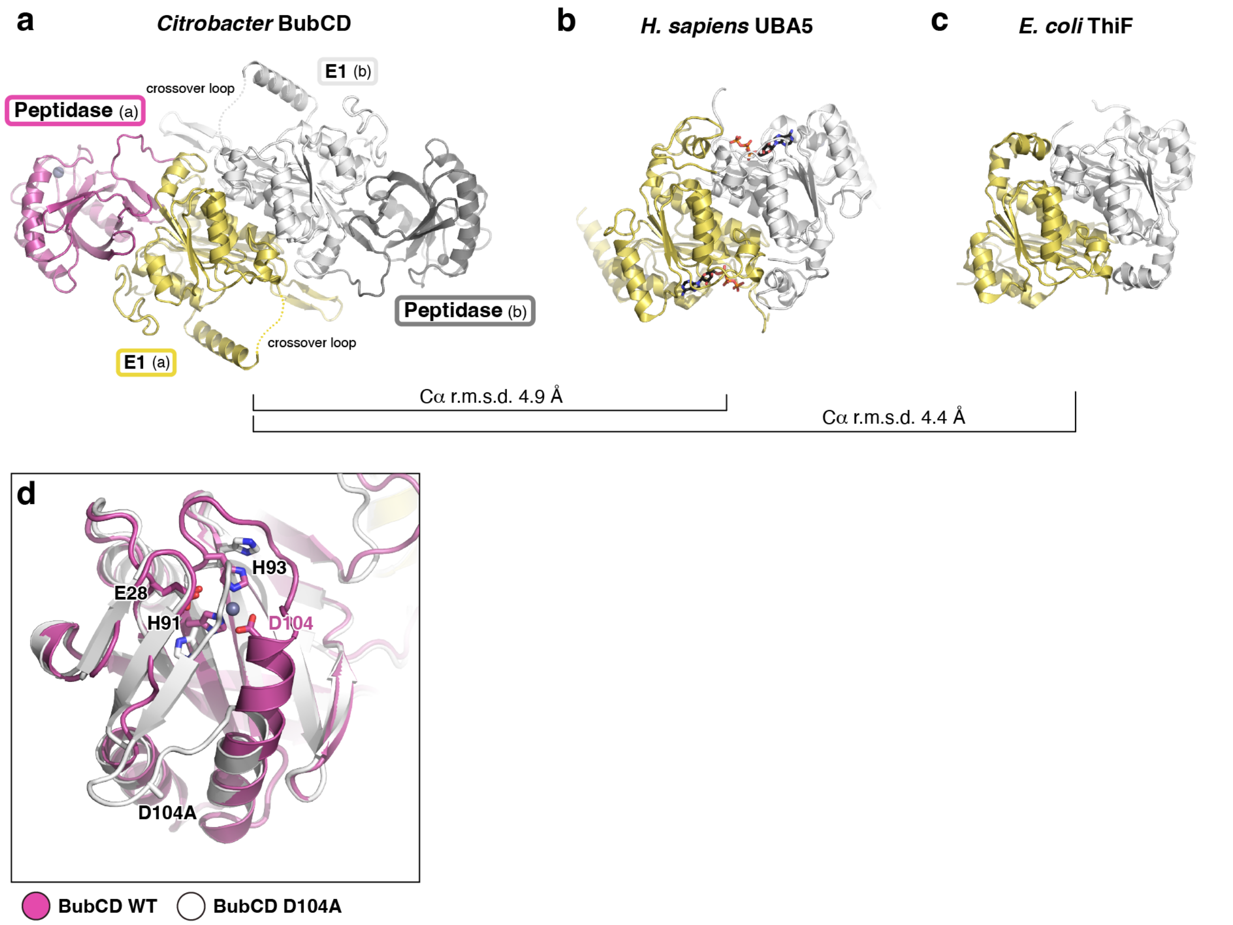

**Figure S2. Structure of *Citrobacter* BubCD. (a)** Structure of the *Citrobacter* BubCD homodimer, with protomer (a) colored as in **Figure 2a** and protomer (b) colored gray/white. **(b)** Structure of the *H. sapiens* homodimeric E1 protein UBA5 (PDB ID 6H78)[18], with one protomer colored yellow and the second protomer colored white. Bound ATP molecules are shown as sticks. **(c)** Structure of the *E. coli* ThiF homodimer (PDB ID 1ZUD)[40], with one protomer colored yellow and the second protomer colored white. **(d)** Overlay of the BubCD N-terminal peptidase domain in the structure of wild-type BubCD (purple) and BubCD$^{D104A}$ (white). Residues 93-108 show a structural rearrangement in BubCD$^{D104A}$, resulting in destruction of the active site and BubA binding site.

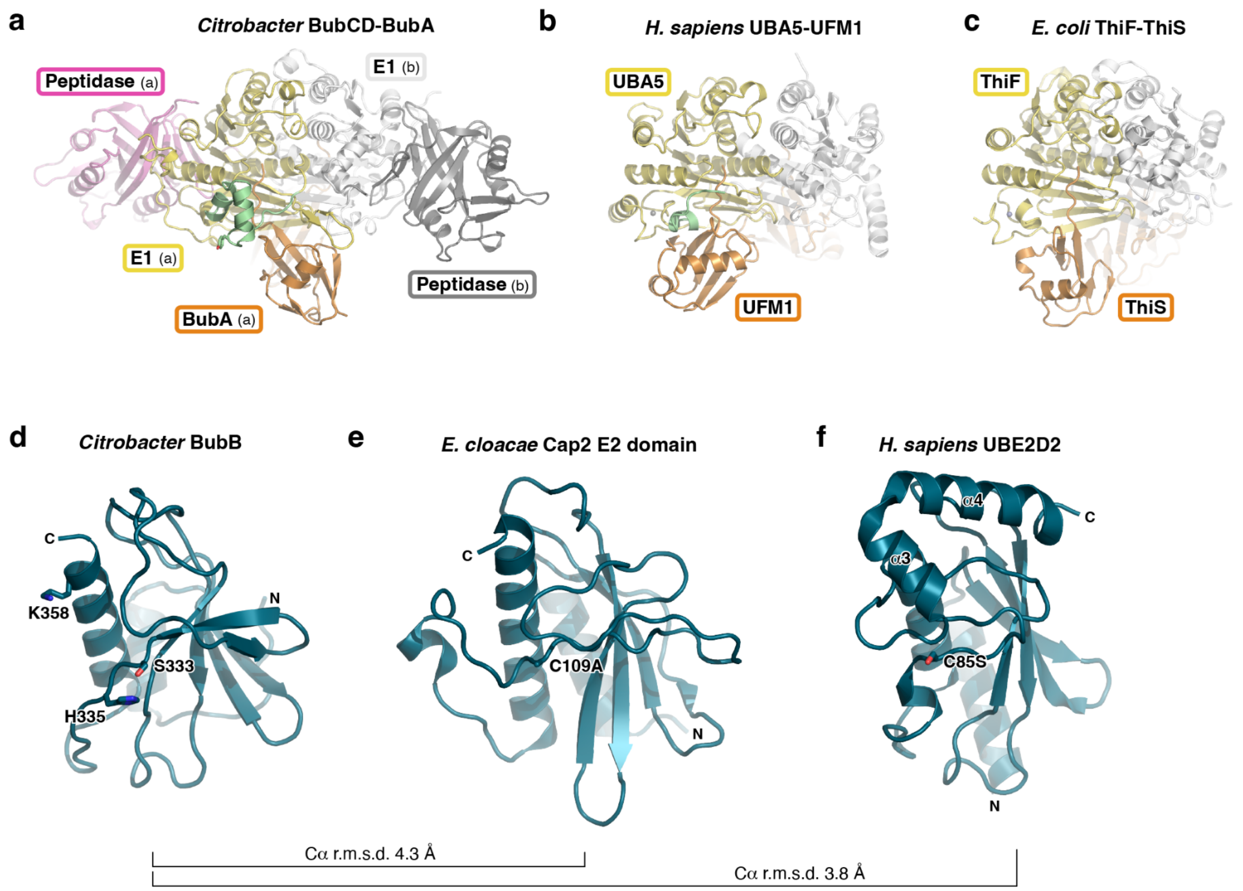

**Figure S3. Structures of *Citrobacter* BubCD bound to BubA and BubB (a)** Structure of the *Citrobacter* BubCD$^{D104}$-BubA$^{155-229}$ complex, with one BubCDD104A protomer colored as in Figure 2a and the second protomer colored gray/white, and BubA colored orange. **(b)** Structure of human UBA5-UFM1 (PDB ID 5IAA)[41], with one UBA5 protomer colored yellow and the second protomer colored white, and UFM1 colored orange.The crossover loop of one protomer is colored light green. **(c)** Structure of *E. coli* ThiF-ThiS (PDB ID 1ZUD)[40], with one ThiF protomer colored yellow and the second protomer colored white, and ThiS colored orange. **(d)** Structure of *Citrobacter* BubB, with putative active site residues S333 and H335 shown as sticks. Lysine K358, mutation of which eliminates formation of the BubA-BubB product in *in vitro* reactions, is also shown as sticks. **(e)** Structure of the E2 domain of *E. cloacae* Cap2 (PDB ID 7TO3)[2], with catalytic cysteine C109 (mutated to alanine) shown as sticks. **(f)** Structure of *H. sapiens* UBE2D2 (PDB ID 4DDG)[42], with catalytic serine C85 (mutated to serine) shown as sticks.

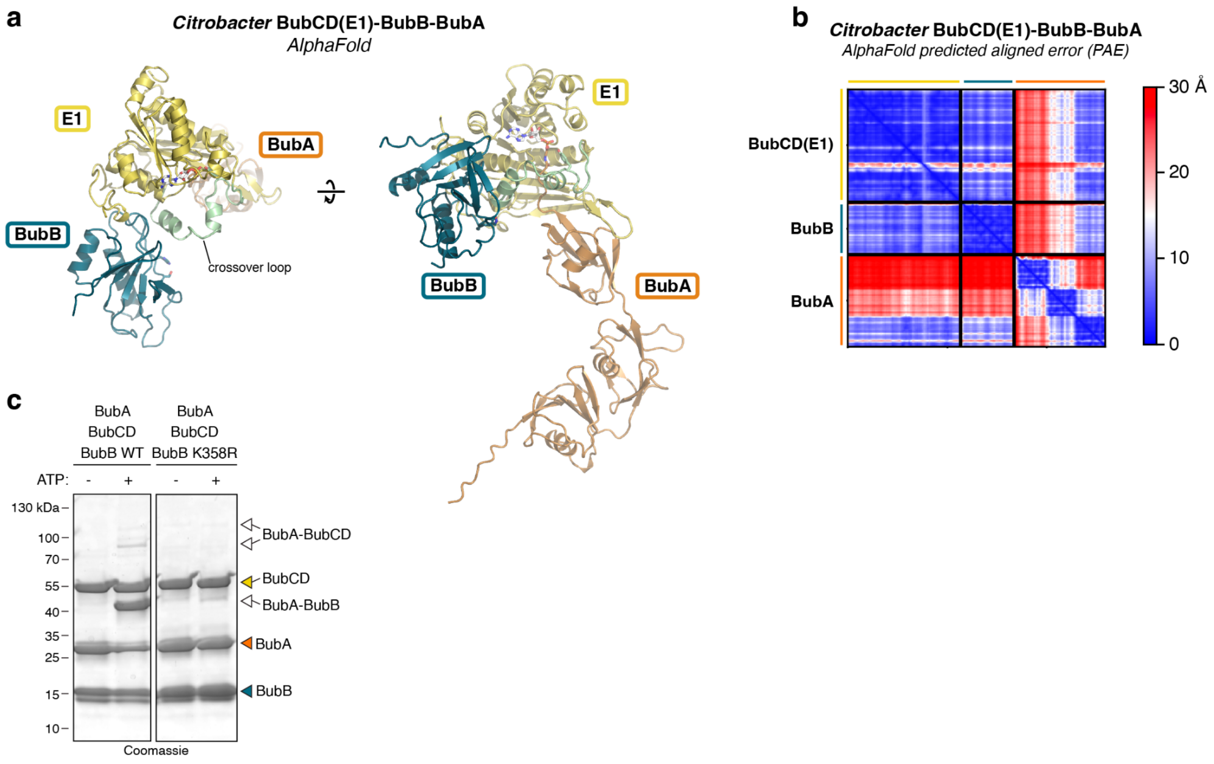

**Figure S4. Type I Bub proteins function through oxyester intermediates. (a)** Two views of an AlphaFold 2-predicted structure of *Citrobacter* BubCD (E1 domain only) bound to BubB and full-length BubA. Domains are colored as in **Figure 3a**. **(b)** AlphaFold 2 predicted aligned error (PAE) plot for the structure shown in panel (a). **(c)** SDS-PAGE analysis of *in vitro* ubiquitination reactions with *Citrobacter* Bub proteins, showing that the BubA-BubB product does not form when BubB's sole lysine residue is mutated to arginine (K358R).

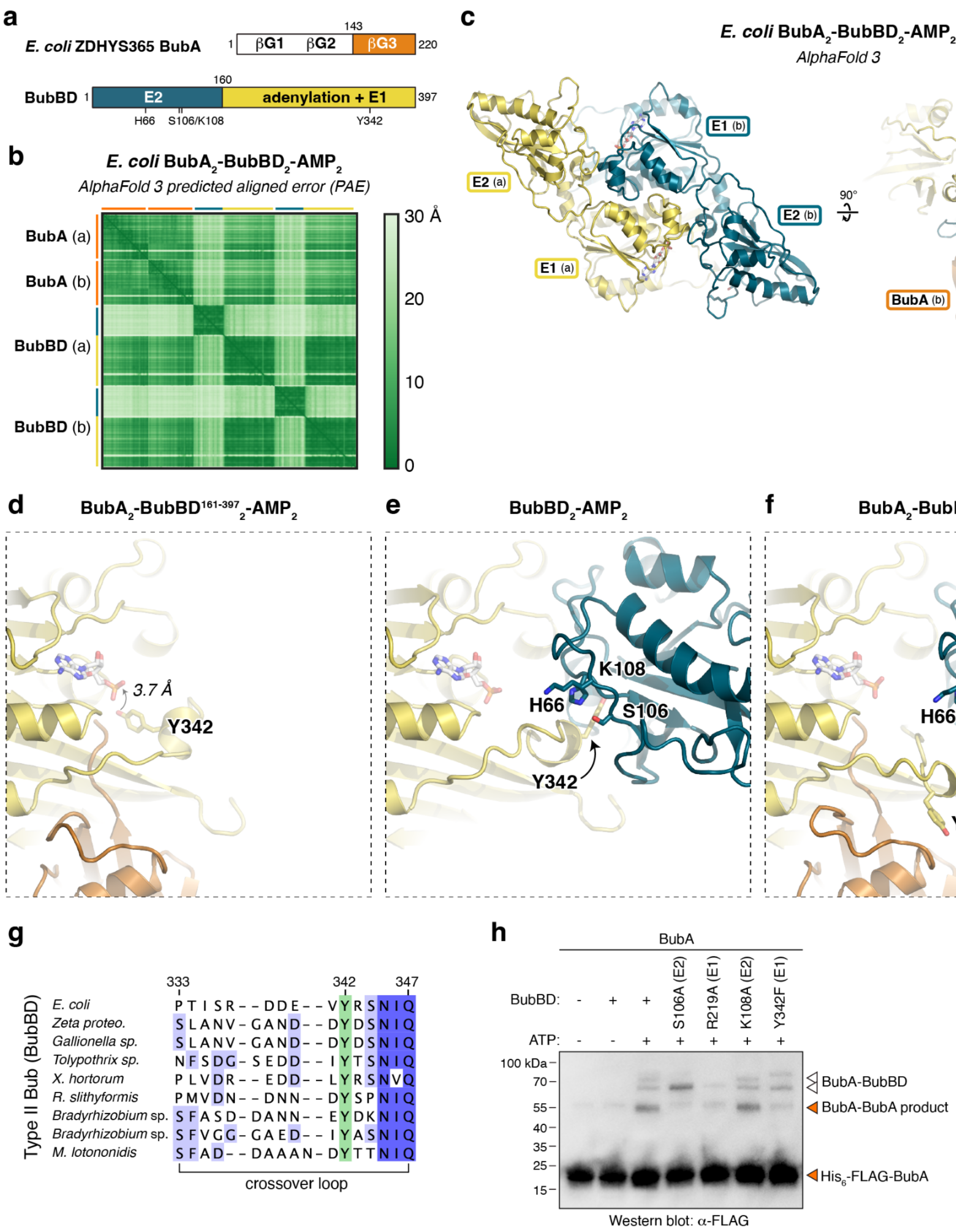

**Figure S5. Type II Bub proteins function through oxyester intermediates. (a)** Domain schematics of *E. coli* ZDHYS365 BubA and BubBD (see **Table S2**). **(b)** AlphaFold 3 predicted aligned error (PAE) plot for a prediction of *E. coli* BubA (two copies), BubBD (two copies), and adenosine monophosphate (AMP; two molecules). **(c)** Two views of the predicted structure of *E. coli* $BubA_2$-$BubBD_2$-$AMP_2$. Domains are colored as in panel (a), and only the C-terminal β-grasp domain of BubA is shown. **(d-f)** Closeup views of predicted BubBD catalytic sites in AlphaFold 3 predictions of *E. coli* $BubA_2$-$BubBD^{161\text{-}397}{}_2$-$AMP_2$ (d), $BubBD_2$-$AMP_2$ (e), and $BubA_2$-$BubBD_2$-$AMP_2$ (f). Domains are colored as in panel (a), and predicted catalytic residues are shown as sticks and labeled. **(g)** Sequence alignment of the crossover loop in Type II Bub BubBD proteins (from an alignment of 400 unique BubBD proteins, see **Table S1**). Proteins shown are from *E. coli* ZDHYS365 (NCBI WP_063617865.1; numbering shown), Zeta proteobacterium SCGC AB-

604-B04 (IMG 2264879632), *Gallionella* sp. SCGC AAA018-N21 (IMG 2264885673), *Tolypothrix* sp. PCC 7601 (IMG 2501546803), *Xanthomonas hortorum pv. gardneri* 7 (IMG 2501866950), *Runella slithyformis* LSU4, DSM 19594 (IMG 2505789128), *Bradyrhizobium* sp. WSM1417 (IMG 2507506257), *Bradyrhizobium* sp. WSM471 (IMG 2508545363), and *Microvirga lotononidis* WSM3557 (IMG 2509076069). **(h)** *In vitro* reconstitution of Ubl conjugation activity of *E. coli* Bub operon proteins. Mutation of BubBD S106, R219, and Y342 all disrupt formation of the likely BubA-BubA product, while mutation of BubBD K108 does not.

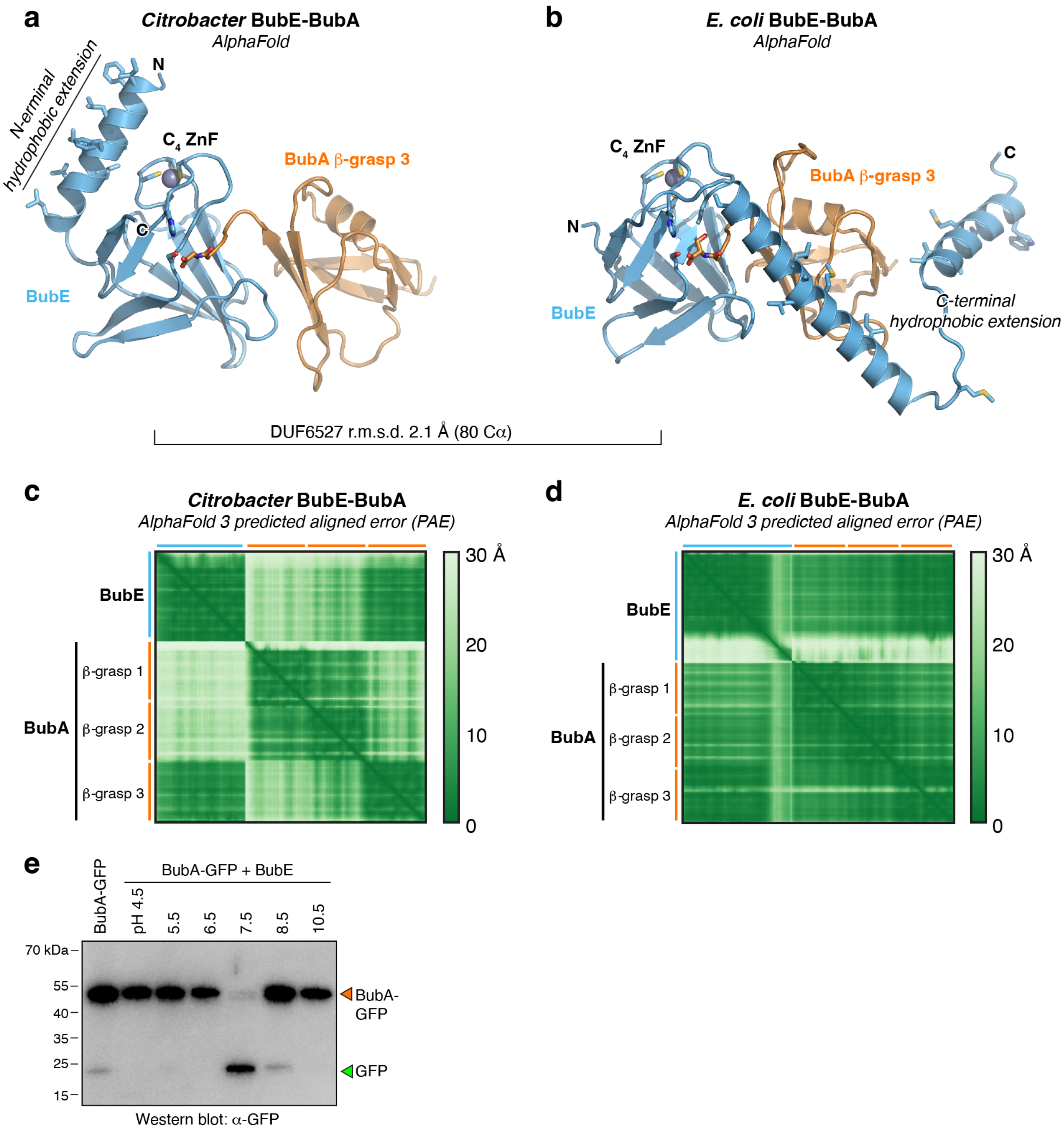

**Figure S6. BubE/DUF6527 is a serine protease. (a)** AlphaFold 3-predicted structure of *Citrobacter* BubE (blue) bound to BubA (orange; only the third β-grasp domain shown) and $Zn^{2+}$ (gray sphere). The N-terminal hydrophobic extension of BubE is indicated, and the residues involved in $Zn^{2+}$ binding (C57, C59, C61, and C99) and the putative catalytic dyad (S92 and H102) are shown as sticks. **(b)** AlphaFold 3-predicted structure of *E. coli* ZDHYS365 BubE (blue) bound to BubA (orange; only the third β-grasp domain shown) and $Zn^{2+}$ (gray sphere). The C-terminal hydrophobic extension of BubE is indicated, and the residues involved in $Zn^{2+}$ binding (C34, C36, C38, and C73) and the putative catalytic dyad (S65 and H76) are shown as sticks. *Citrobacter* and *E. coli* BubEs' DUF6527 domains overlay with an r.m.s.d. of 2.1 Å over

80 Cα atoms. **(c)** AlphaFold 3 predicted aligned error (PAE) plot for the structure prediction of Citrobacter BubE-BubA shown in panel (a). **(d)** AlphaFold 3 predicted aligned error (PAE) plot for the structure prediction of *Citrobacter* BubE-BubA shown in panel (b). **(e)** *In vitro* cleavage of BubA-GFP fusion by *Citrobacter* BubE in buffers ranging from pH 4.5 to 10.5. Proteins are detected by anti-GFP western blot.

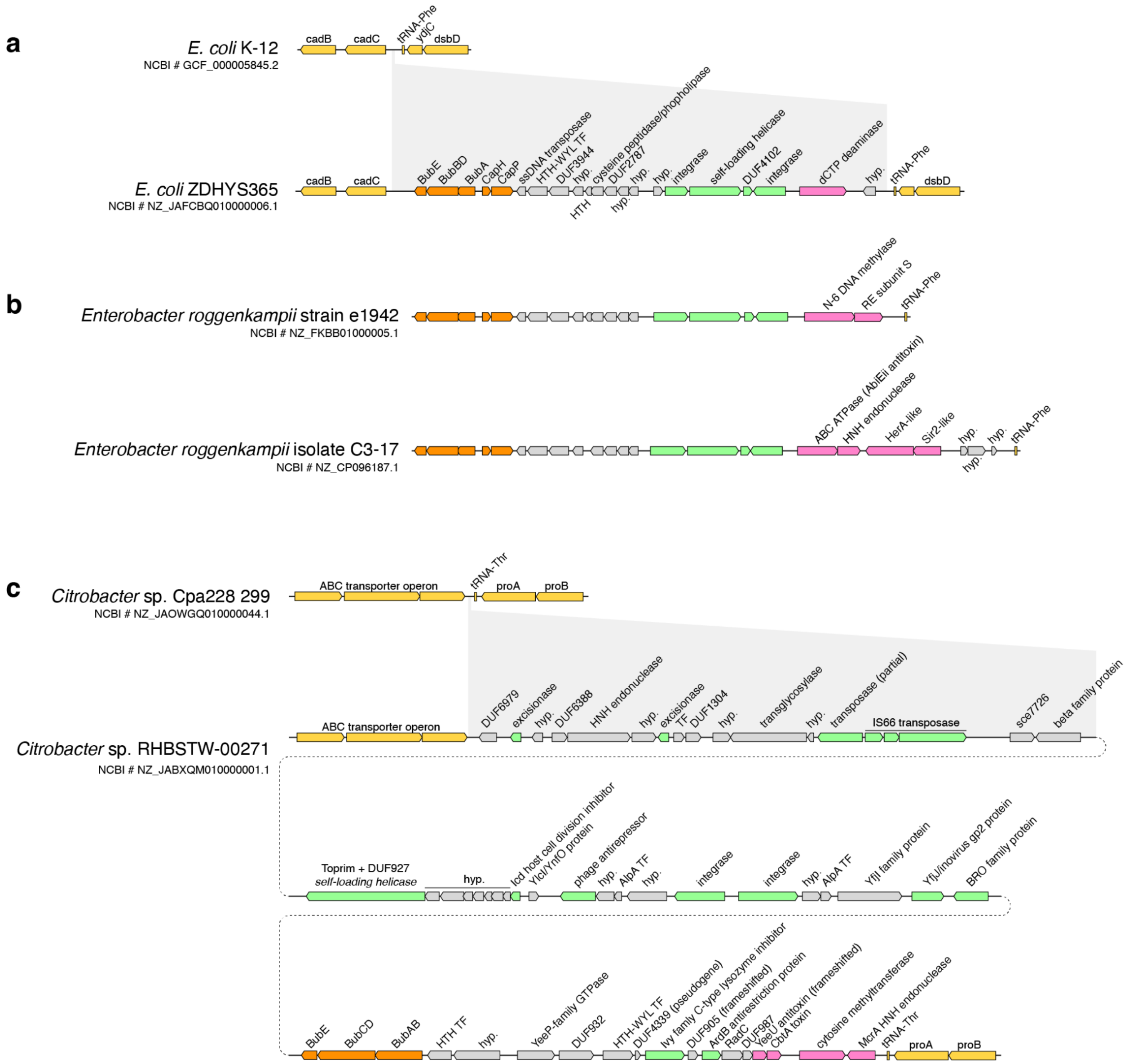

**Figure S7. Genomic context of Bub operons. (a)** Schematic of the equivalent regions of *E. coli* K-12 and *E. coli* ZDHYS365, showing a mobile element inserted near a phenylalanine tRNA gene (tRNA-Phe; yellow) in *E. coli* ZDHYS365. Bub operon is shown in orange, other likely immune systems are shown in pink, and likely replication/integration genes are shown in green. **(b)** Two mobile elements from strains of *Enterobacter roggenkampii*, showing the same overall organization as the *E. coli* ZDHYS365 mobile element but with different immune systems. **(c)** Schematic of the equivalent regions of *Citrobacter* sp. Cpa228_299 and *Citrobacter* sp. RHBSTX-00271, showing a ~45 kb mobile element/prophage locus in *Citrobacter* sp. RHBSTX-00271 inserted near a threonine tRNA gene (tRNA-Thr; yellow). Bub operon is shown in orange, other likely immune systems are shown in pink, and likely replication/integration genes plus other characteristic phage genes are shown in green.
